# GNS561, a clinical-stage PPT1 inhibitor, is efficient against hepatocellular carcinoma via modulation of lysosomal functions

**DOI:** 10.1101/2020.09.30.320010

**Authors:** Sonia Brun, Eric Raymond, Firas Bassissi, Zuzana Macek Jilkova, Soraya Mezouar, Madani Rachid, Marie Novello, Jennifer Tracz, Ahmed Hamaï, Gilles Lalmanach, Lise Vanderlynden, Eloïne Bestion, Raphael Legouffe, Jonathan Stauber, Thomas Schubert, Maximilian G. Plach, Jérôme Courcambeck, Cyrille Drouot, Guillaume Jacquemot, Cindy Serdjebi, Gael Roth, Jean-Pierre Baudoin, Christelle Ansaldi, Thomas Decaens, Philippe Halfon

**Affiliations:** Genoscience Pharma, Marseille, France; Medical Oncology, Paris Saint-Joseph Hospital, Paris, France; Institute for Advanced Biosciences, Research Center UGA / Inserm U 1209 / CNRS 5309, La Tronche, France; University of Grenoble Alpes, Faculté de médecine, France; Clinique Universitaire d’Hépato gastroentérologie, Pôle Digidune, CHU Grenoble, France; Institut Necker-Enfants Malades, Inserm U1151-CNRS UMR 8253, Paris, France; University of Paris Descartes-Sorbonne Paris Cité, Paris, France; INSERM, UMR1100, Centre d’Etude des Pathologies Respiratoires, Equipe “Mécanismes Protéolytiques dans l’Inflammation“, Tours, France; University of Tours, Tours, France; CNRS, IRD, MEPHI, IHU Méditerranée Infection, University of Aix Marseille, Marseille, France; ImaBiotech, Loos, France; ImaBiotech, Billerica, USA; 2Bind GmbH, Regensburg, Germany; University of Aix Marseille, IRD, APHM, MEPHI, IHU Méditerranée Infection, Marseille, France

**Keywords:** liver cancer, lysosome, autophagy, antitumor, mTOR

## Abstract

**Background & Aims:** Hepatocellular carcinoma (HCC) is the most frequent primary liver cancer. Autophagy inhibitors have been extensively studied in cancer but, to date, none has reached efficacy in clinical trials.

**Approach & Results:** To explore the antitumor effects of GNS561, a new autophagy inhibitor, we first achieved in vitro assays using various human cancer cell lines. Having demonstrated that GNS561 displayed high liver tropism using mass spectrometry imaging, the potency of GNS561 on tumor was evaluated in vivo in two HCC models (human orthotopic patient-derived xenograft mouse model and diethylnitrosanime-induced cirrhotic immunocompetent rat model). Oral administration of GNS561 was well tolerated and decreased tumor growth in these two models. GNS561 mechanism of action was assessed in an HCC cell line, HepG2. We showed that due to its lysosomotropic properties, GNS561 could reach and inhibited its enzyme target, palmitoyl-protein thioesterase 1, resulting in lysosomal unbound Zn^2+^ accumulation, impairment of cathepsin activity, blockage of autophagic flux, altered location of mTOR, lysosomal membrane permeabilization, caspase activation and cell death.

**Conclusions:** GNS561, currently tested in a global Phase 1b/2a clinical trial against primary liver cancer, represents a promising new drug candidate and a hopeful therapeutic strategy in cancer treatment.

With an estimated 782,000 deaths in 2018, hepatocellular carcinoma (HCC) stands as the most common primary liver cancer and constitutes the fourth leading cause of cancer-related death worldwide (1). The rising incidence of HCC, the high worldwide mortality rate, and limited therapeutic options at advanced stages, make HCC a significant unmet medical need.

Autophagy-related lysosomal cell death, either alone or in connection with several other cell death pathways, has been recognized as a major target for cancer therapy (2). Dysregulated autophagic-lysosomal activity and mTOR signaling were shown to allow cancer cells to become resistant to the cellular stress induced by chemotherapy and targeted therapy (3). Recently, several lysosome-specific inhibitors were shown to target palmitoyl-protein thioesterase 1 (PPT1), resulting in the modulation of protein palmitoylation and autophagy, and antitumor activity in melanoma and colon cancer models (4, 5).

Chloroquine (CQ) and hydroxychloroquine (HCQ) have been used for more than 50 years to prevent and treat malarial infections and autoimmune diseases. Based on the lysosomotropic properties and the capacity for autophagy inhibition, these molecules have been proposed as active drugs in cancer (6–9). Over 40 clinical trials have been reported to evaluate the activity of both CQ or HCQ as single agent or in combination with chemotherapy in several tumor types (6–8. However, the required drug concentrations to inhibit autophagy were not achieved in humans, leading to inconsistent results in cancer clinical trials (5, 10). This prompted research to identify novel compounds with potent inhibitory properties against autophagy for cancer therapy.

We previously reported that GNS561 was efficient in intrahepatic cholangiocarcinoma (iCCA) by inhibiting late-stage autophagy (11). In this study, we investigated the mechanism of action of GNS561. We identified lysosomal PPT1 as a target of GNS561. Exposure to GNS561 induced lysosomal accumulation of unbound zinc ion (Zn^2+^), inhibition of PPT1 and cathepsin activity, blockage of autophagic flux and mTOR displacement. Interestingly, these effects resulted in lysosomal membrane permeabilization (LMP) and caspase activation that led to cancer cell death. This mechanism was associated with dose-dependent inhibition of cancer cell proliferation and tumor growth inhibition in two HCC in vivo models. These data establish PPT1 and lysosomes as major targets for cancer cells and led to the development of a clinical program investigating the effects of GNS561 in patients with advanced HCC.

## Materials and Methods

### Materials

Information on the materials used in this study is provided in the Supplementary Material online.

### Animal treatment

The protocol for the animal experiments was approved by animal Ethics Committees and followed Guidelines on the Humane Treatment of Laboratory Animals. Detailed information is provided in the Supplementary Material online.

## Results

### GNS561 displays activity against human cancer cell lines and patient-derived cells

The effects of GNS561 on cell viability were investigated in a panel of human cancer cell lines, including HCC, iCCA and colon, renal cell, breast, prostate, lung, and ovarian carcinoma as well as acute myeloid leukemia, glioblastoma, and melanoma. As shown in Table 1, GNS561 showed potent antitumor activity ranging from 0.22 ± 0.06 μM for the most sensitive cell line (LN-18, a glioblastoma cell line) to 7.27 ± 1.71 μM for the least sensitive cell line (NIH:OVCAR3, an ovarian cancer cell line). GNS561 was at least 10-fold more effective than HCQ in cultured cancer cells. GNS561 also displayed activity in primary HCC patient-derived cells and was on average 3-fold more potent than sorafenib, a reference drug in HCC treatment (mean IC_50_ 3.37 ± 2.40 μM for GNS561 vs 10.43 ± 4.09 μM for sorafenib).

**Table 1.** In vitro activity of GNS561 and HCQ in human cancer cell lines (left, IC_50_ ± SD, μM) and in vitro activity of GNS561 and sorafenib in primary hepatocellular carcinoma (HCC) patient-derived cells (right, IC_50_, μM).

|  |  | Mean IC<sub>50</sub> ± SD (μM) |  |  | IC<sub>50</sub> (μM) |  |
| --- | --- | --- | --- | --- | --- | --- |
| Cancer type | Cell lines | GNS561 | HCQ | Primary HCC patient-derived cells | GNS561 | sorafenib |
| Colon Carcinoma | HCT-116 | 1.22 ± 0.15 | 14.41 ± 1.5 | LI0050 | 3.54 | 9.12 |
|  | HT-29 | 1.35 ± 0.04 | 24.18 ± 5.14 | LI0574 | 2.41 | 8.65 |
| Renal Cell Carcinoma | 786-O | 1.72 ± 0.17 | 21.65 ± 3.15 | LI0612 | 6.93 | 17.94 |
|  | CAKI-1 | 1.10 ± 0.19 | 17.69 ± 1.29 | LI0752 | 0.49 | 6.34 |
| Ovarian Cancer | NIH:OVCAR3 | 7.27 ± 1.71 | 98.01 ± 12.75 | LI0801 | 2.07 | 5.7 |
| Melanoma | A375 | 1.2 ± 0.13 | 12.27 ± 2.8 | LI1005 | 3.16 | 14.49 |
|  | SK-MEL-28 | 1.81 ± 0.5 | 22.78 ± 2.65 | LI1098 | 6.95 | 10.85 |
| Breast Cancer | MDA-MB-231 | 2.17 ± 0.14 | 14.13 ± 3.06 | LI1646 | 1.44 | 10.33 |
| Prostate Cancer | DU-145 | 1.09 ± 0.18 | 45.74 ± 0.55 | Mean | 3.37 ± 2.40 | 10.43 ± 4.09 |
|  | PC-3 | 2.56 ± 0.23 | 43.43 ± 6.04 |  |  |  |
| Lung Cancer | A549 | 1.69 ± 0.34 | 14.33 ± 1.59 |  |  |  |
|  | NCI-H358 | 2.54 ± 0.34 | 54.07 ± 14.19 |  |  |  |
| HCC | HepG2 | 0.47 ± 0.15 | 11.55 ± 1.52 |  |  |  |
|  | Huh7 | 0.88 ± 0.31 | 13.62 ± 0.71 |  |  |  |
| Glioblastoma | LN-229 | 0.60 ± 0.24 | 10.87 ± 1.23 |  |  |  |
|  | LN-18 | 0.22 ± 0.06 | 5.27 ± 0.74 |  |  |  |
| Acute Myeloid Leukemia | KG-1 | 5.86 ± 1.64 | 43.92 ± 2.76 |  |  |  |
|  | Mean | 1.99 ± 1.86 | 27.52 ± 23.28 |  |  |  |

### GNS561 has antitumor properties in HCC in vivo models

The whole-body tissue distribution of GNS561 was investigated in Sprague Dawley rats after repeated oral administration of GNS561 at a dose of 40 mg/kg/day for 28 days. Seven hours after the last administration, the GNS561 level was measured by mass spectrometry imaging in the liver, lung, stomach, brain, eye, salivary gland, kidney, heart, fat, muscle, testis and skin (Fig. 1A). GNS561 mainly accumulated in the liver, stomach and lung as shown by the calculated organ/blood ratio (Fig. 1B). Lower concentrations of GNS561 were also detected in eyes, skin, brain and testis, indicating that GNS561 crosses the blood/brain barrier and the blood/testis barrier to a limited extent (brain to blood and testis to blood ratios were 0.21 and 0.40, respectively). High liver concentrations were also highlighted in rats after repeated administration of GNS561 at the dose level of 15 and 30 mg/kg/day for 28 days and 50 mg/kg/day for 21 days (data not shown).

**Fig. 1.**
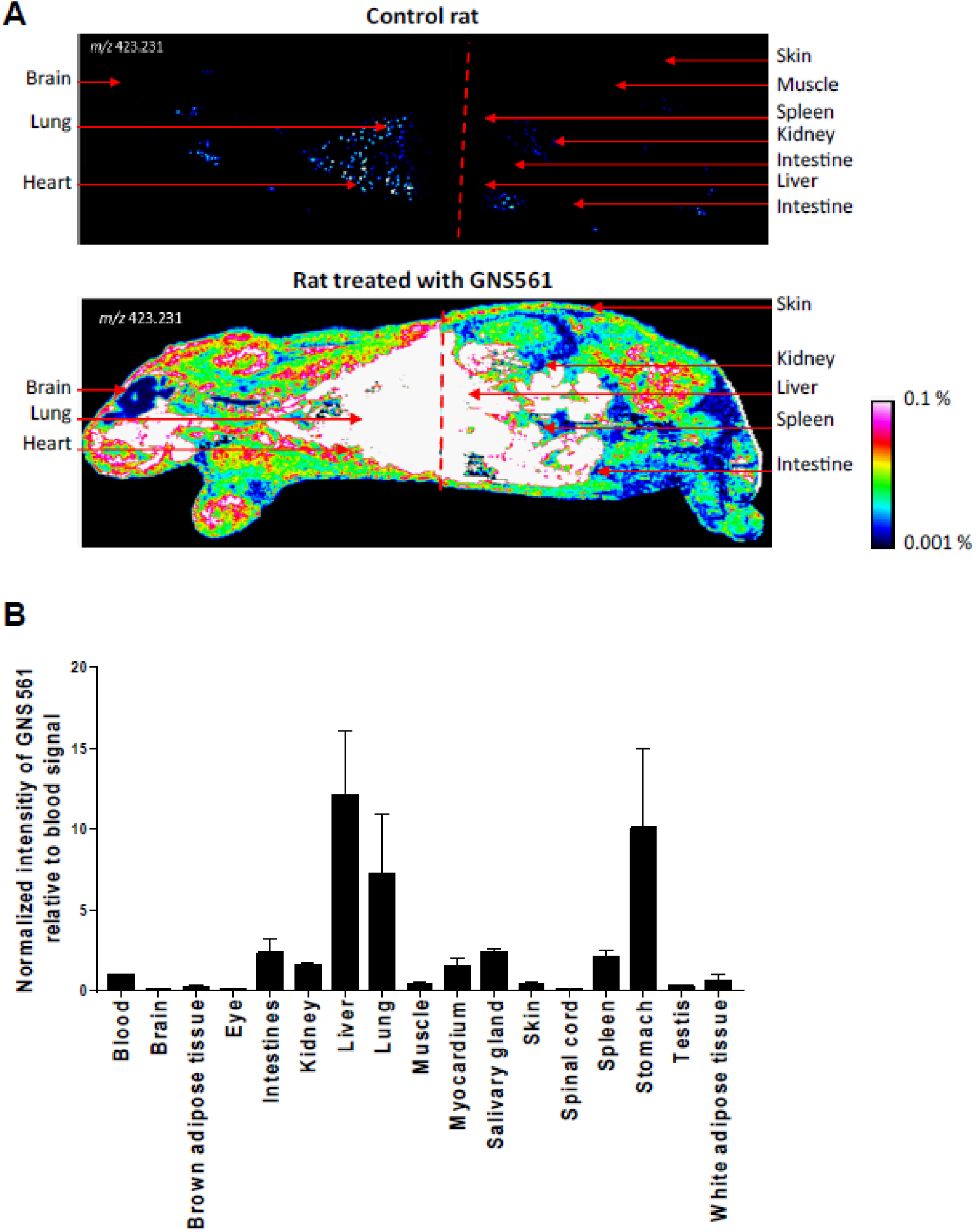
Whole body tissue distribution of GNS561. (A) Mass spectrometry imaging of a control rat (top) and a rat treated with GNS561 at a dose of 40 mg/kg/day for 28 days (bottom). (B) Normalized intensity of GNS561 relative to blood signal in several organs of GNS561-treated rats (Mean + SEM, n=2 except for eye, n=1). Of note, the GNS561 liver-to-blood ratio is underestimated due to liver signal saturation.

Based on the high concentrations of GNS561 in the liver and potent in vitro activity against HCC cells, the effects of GNS561 were investigated in vivo using two liver cancer models, including the human HCC orthotopic patient-derived LI0752 xenograft mouse model and the diethylnitrosanime (DEN)-induced cirrhotic immunocompetent rat model of HCC.

In the HCC patient-derived LI0752 xenograft BALB/c nude mouse model, tumor volume and weight were reduced by 37.1% and 34.4%, respectively, in mice treated with GNS561 at 50 mg/kg compared to the control (Supplementary Fig. 1A and B). Consistently, GNS561 treatment induced a decrease in serum AFP levels in a dose-dependent manner and was significantly different from the control at days 21 and 28 after treatment (Supplementary Fig. 1C-G).

Since HCC often develops in cirrhotic livers in humans, we further characterized the antitumor effects of GNS561 in a DEN-induced cirrhotic rat model of HCC. Rats with already developed HCC were either treated with sorafenib at 10 mg/kg, GNS561 at 15 mg/kg, or the combination of both drugs (Supplementary Fig. 2). In this model, tumor progression was significantly reduced by sorafenib (33.0%) and GNS561 (33.0%) compared to an untreated control group, and the greatest decrease in tumor progression was observed by the combination (68%) that displayed an additive effect (Fig. 2A). Magnetic resonance imaging analyses further showed a significant increase in the mean tumor size of 9.97 ± 0.97 mm in control rats compared to 6.45 ± 0.35 mm with sorafenib, 5.48 ± 1.00 mm in GNS561 and 3.83 ± 0.52 mm in the combination group (Fig. 2B). Following liver resection, the macroscopic counting of tumor nodules revealed significantly lower numbers in all treated groups compared to the control group (Fig. 2C). Immunohistochemical analyses of liver tumors showed a significantly lower Cyclin D1-positive nuclear staining in the tumors of rats treated with GNS561 or by the combination of GNS561 with sorafenib compared to the control group (Fig. 2D). GNS561 and combination treatments also significantly reduced Ki67 staining compared to the control group (Fig. 2E). The effects on Cyclin D1 and Ki67 were primarily related to GNS561 exposure, as sorafenib alone showed no statistically significant differences in Cyclin D1 or Ki67 staining compared to the control group. Our results further showed that GNS561 and the combination treatment did not interfere with lipid or glucose metabolism or kidney function but slightly affected some liver functions (Table S1).

**Fig. 2.**
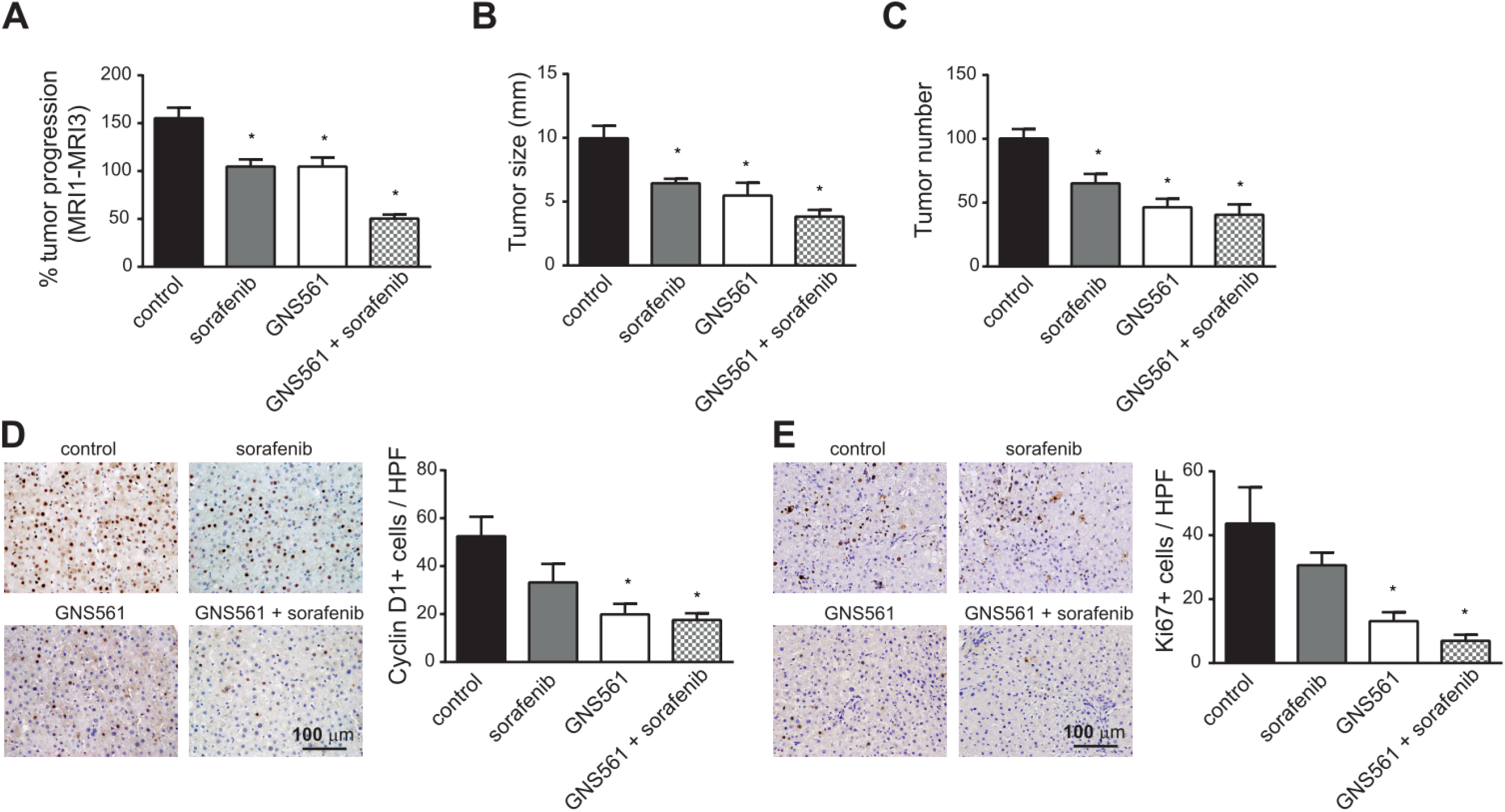
GNS561 activity in a diethylnitrosanime-induced cirrhotic rat model of hepatocellular carcinoma. (A) Tumor progression assessment by comparison of tumor size obtained by magnetic resonance imaging (MRI) 1 and MRI 3 in the control, sorafenib at 10 mg/kg, GNS561 at 15 mg/kg and combination (GNS561 + sorafenib) groups. Macroscopic examination of livers with assessments of (B) tumor size and (C) tumor number at the surface of livers. (D) Representative images of nuclear Cyclin D1 staining and quantification of Cyclin D1-positive staining per high-power field (HPF). (E) Representative images of nuclear Ki67-positive staining and quantification of Ki67 staining per HPF. For all studies, mice n ≥ 6 per group. Data represent the mean + SEM. Comparison of means was performed by one-way ANOVA with Dunnett’s post hoc analysis. * represents significant difference, at least p < 0.05.

### GNS561 activates the caspase-dependent apoptosis pathway

We further wanted to characterize the antitumor effect of GNS561 and to determine whether GNS561 could trigger apoptotic cell death. To this end, annexin V/propidium iodide (PI) analysis was performed by flow cytometry after 48 h of GNS561 exposure in HepG2 cells. Early (Annexin V+/PI-staining) and late (Annexin V+/PI- staining) apoptosis increased in a dose-dependent manner after GNS561 exposure (Fig. 3A). The induction of apoptosis was confirmed by immunodetection of poly-ADP-ribose polymerase cleavage in GNS561-treated cells (Fig. 3B). We further examined whether GNS561-induced apoptosis was related to caspase activation. Cleaved caspase 3 was detected using immunoblot analysis (Fig. 3C). The induction of caspase activation was confirmed by flow cytometry (Fig. 3D) and luminescence analysis (Fig. 3E). After 6 h of exposure, GNS561 had no effect on caspase 8 and caspase 3/7 activity in HepG2 cells (Fig. 3E). In contrast, activation of caspase 8 and caspase 3/7 was observed after 24 h of treatment with GNS561, and this effect was sustained at 30 h. A decrease in cell viability was concomitant with caspase activation (Fig. 3E). Moreover, to confirm that GNS561-induced cell death is caspase-dependent apoptosis, pretreatment (1 h) with the cell-permeable pan-caspase inhibitor Z-VAD-FMK (5 μM) was performed. Cell viability was restored in the presence of Z-VAD-FMK (Fig. 3F), further confirming that GNS561 induced a caspase-dependent apoptotic cell death.

**Fig. 3.**
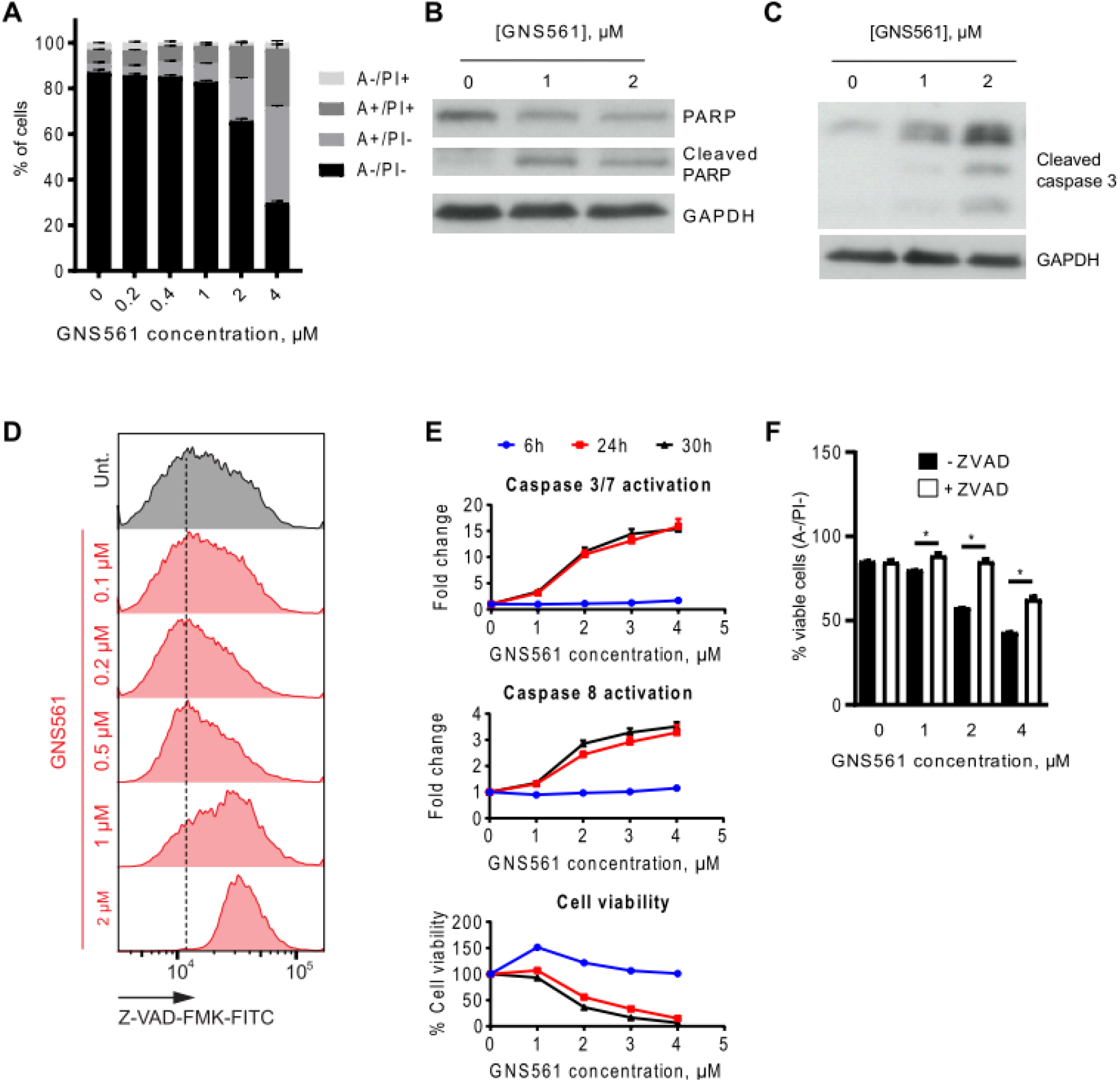
GNS561 induces apoptotic cell death in HepG2 cells in a dose and time-dependent manner through caspase activation. (A) Annexin V (A)/propidium iodide (PI) analysis by flow cytometry after 48 h of GNS561 treatment. (B) Representative immunoblotting of the cleaved and non-cleaved forms of poly-ADP-ribose polymer (PARP) after 24 h of GNS561 treatment. (C) Representative immunoblotting of cleaved caspase 3 levels after 24 h of treatment with GNS561. (D) Caspase-glow analysis by flow cytometry after 48 h of treatment with GNS561. (E) Cell viability and activation of caspase 3/7 and 8 after 6, 24 and 30 h of treatment with GNS561. (F) Viable cell (A-/PI-) analysis by flow cytometry after pretreatment with Z-VAD-FMK (ZVAD) at 5 μM for 1 h and then treatment with ZVAD at 5 μM and GNS561 for 48 h. For all blots, glyceraldehyde-3-phosphate dehydrogenase (GAPDH) was used as a loading control. For all studies, n ≥ 3 biological replicates. Data represent the mean + SEM. For comparison, Student t-test was used. * represents significant difference, at least p < 0.05.

### GNS561 is a lysosomotropic agent

The intracellular localization of GNS561 in HepG2 cells was visualized using GNS561D, the photoactivable analog of GNS561 containing a diazide moiety (Fig. 4A). GNS561D showed a punctuate fluorescent signal that colocalized with the intracellular vesicle-like structure stained by LAMP1 (Fig. 4B), demonstrating that GN561 accumulated in lysosomes and is a lysosomotropic agent. Pretreatment with NH_4_Cl, a weak base that rapidly increases lysosomal pH, was further used to validate the lysosomotropic character of GNS561. As shown in Fig. 4B, NH_4_Cl pretreatment strongly prevented lysosomal accumulation of GNS561D. Then, we investigated whether GNS561 lysosomotropism was related to induced cell death. For this purpose, HepG2 cells were pretreated for 2 h with NH_4_Cl and then treated with GNS561 for 24 h. Although a concentration of 20 mM NH_4_Cl alone slightly decreased viability (Fig. 4C), it significantly attenuated the larger decrease in viability induced by GNS561. These results were confirmed by pretreatment with bafilomycin A1 (Baf A1), an inhibitor of the vacuolar H+-ATPase (Supplementary Fig. 3). Therefore, disrupting GNS561 lysosomal localization protected against GNS561-mediated cell death. These results suggested that GNS561 antitumor activity in HepG2 cells is caused by its lysosomotropism.

**Fig. 4.**
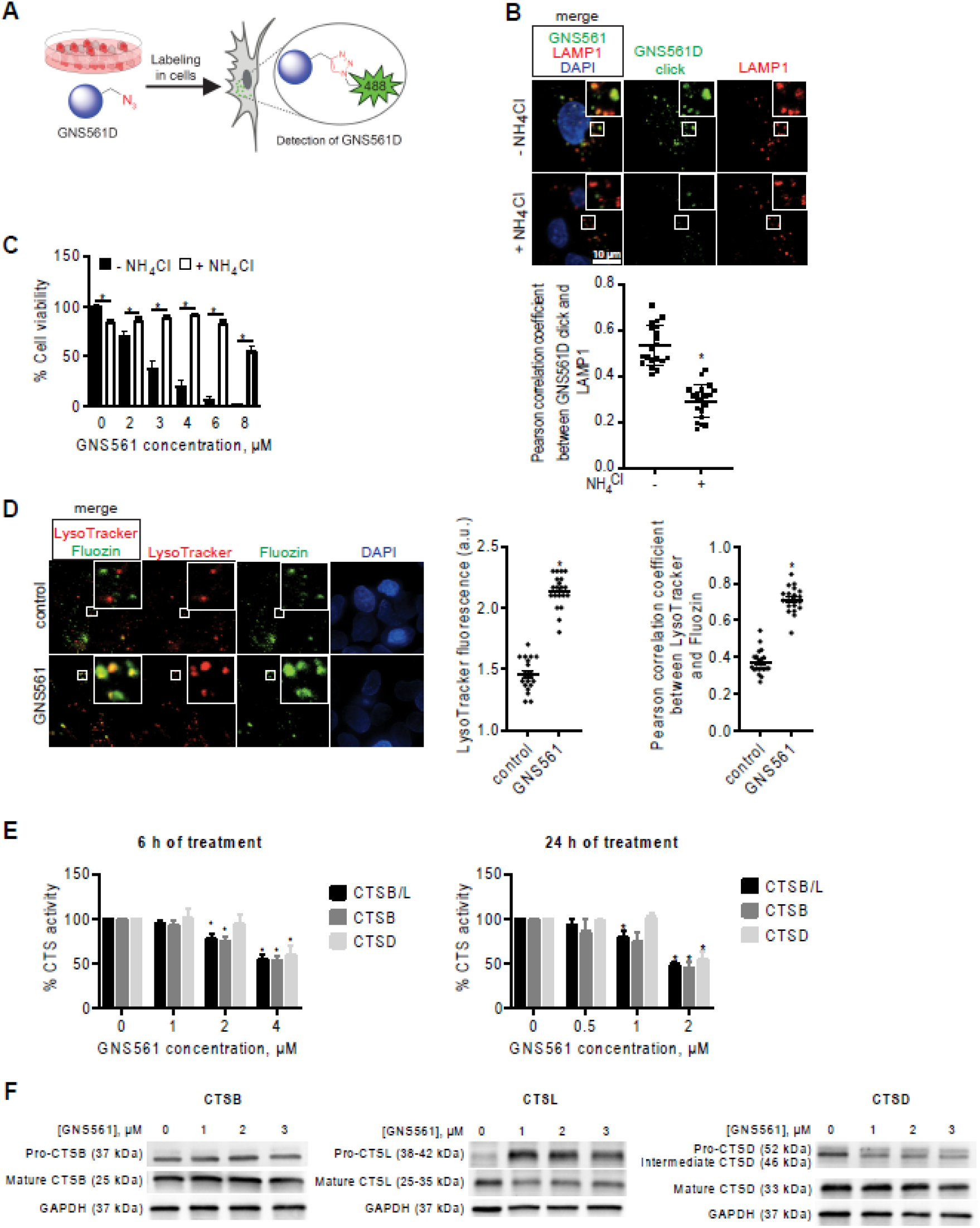
The lysosomotropic agent GNS561 modulates lysosomal functions in the HepG2 cell line. (A) Chemical labeling of GNS561D in cells. (B) Lysosomal localization of GNS561D after NH_4_Cl pretreatment (20 mM) for 30 min and then treatment with GNS561D (10 μM) and NH_4_Cl (20 mM) for 90 min. (C) Cell viability after 24 h of GNS561 exposure in the presence or absence of NH_4_Cl (20 mM). (D) Staining of lysosomes (LysoTracker) and unbound Zn^2+^ (Fluozin) after GNS561 treatment (1 h, 10 μM). Quantification of LysoTracker fluorescence in arbitrary units (a.u.) (middle) and lysosomal unbound Zn^2+^ accumulation by Pearson correlation coefficient between LysoTracker and Fluozin (right). (E) Fold change of peptidase activity of cysteine cathepsins (including both cathepsins B and L) (CTSB/L), cathepsin B (CTSB) and cathepsin D (CTSD) after GNS561 treatment (6 h and 24 h) calculated in comparison with the control condition. (F) Representative immunoblotting of procathepsin B (precursor form) and mature CTSB (left), procathepsin L (precursor form) and mature CTSL (middle) and procathepsin D, intermediate and mature CTSD (right) after GNS561 treatment for 16 h. For all blots, glyceraldehyde-3-phosphate dehydrogenase (GAPDH) was used as a loading control. For all studies, n ≥ 3 biological replicates. Data represent the mean + SEM. For comparison, Student t-test was used for (C), (B) and (D), and one-way ANOVA with Dunnett’s post hoc analysis was performed for (E). * represents significant difference, at least p < 0.05.

### GNS561 modulates lysosomal functions

The GNS561 lysosomotropism-dependent cell death prompted us to examine GNS561 capacity to modulate lysosomal characteristics and functions.

Following continuous exposure to GNS561, staining of LysoTracker, which is a reagent allowing the identification of the lysosomal compartment, increased in HepG2 cells (Fig.4D), suggesting that GNS561 prompted a dose-dependent build-up of enlarged lysosomes. We therefore examined the enzymatic activity of three prominent lysosomal proteinases, two cysteine cathepsins B (CTSB) and L (CTSL), and aspartic cathepsin D (CTSD). After 6 and 24 h of treatment, GNS561 significantly impaired, in a dose-dependent manner, the enzymatic activity of cathepsins (Fig. 4E). However, this decreased activity did not relate to a direct GNS561-dependent inhibition of cathepsin activities (Supplementary Fig. 4). Based on the literature, depressed proteolytic activity of cathepsins may result from an increased Zn^2+^ lysosomal concentration and/or altered maturation of cathepsin precursors. Indeed, it has been described that Zn^2+^ may downregulate the proteolytic activity of CSTB and CTSL (12, 13). We investigated whether GNS561 modified unbound Zn^2+^ localization in HepG2 cells. As shown in Fig. 4D, GNS561 induced a strong accumulation of unbound Zn^2+^ in lysosomes, as evidenced by colocalization of the fluorescent signals of Fluozin and LysoTracker in the merged images. This increase in lysosomal unbound Zn^2+^ could explain the decreased proteolytic activity of CTSL and CTSB. Cathepsins are synthesized as inactive zymogens, which are converted to their mature active forms by other proteases or by autocatalytic processing (14). As depicted in Fig. 4F, GNS561 did not impact CTSB maturation, while it impaired the maturation of both CTSL and CSTD (increase of precursor forms) and decreased their catalytic activity accordingly.

As GNS561 induced lysosomal dysfunction, the effect of GNS561 on the autophagic process was investigated. Herein, we showed that the GNS561-induced accumulation of light chain 3 phosphatidylethanolamine conjugate was not enhanced in the presence of BafA1 (Supplementary Fig. 5), suggesting that GNS561 blocked autophagic flux.

### PPT1 is a target of GNS561

Since PPT1 is critical for lysosomal function and is described to be the molecular target of chloroquine derivatives (4, 5), we investigated whether PPT1 could be a molecular target of GNS561. First, the binding of GNS561 to recombinant PPT1 was analyzed in vitro by nano differential scanning fluorimetry using HCQ as a positive control (4). In the presence of GNS561 and HCQ, we observed a significant dose-dependent decrease in PPT1 melting temperature (Fig. 5A). Additionally, inhibition of PPT1 enzymatic activity was observed in HepG2 cells treated with GNS561 (Fig. 5B). Moreover, the chemical mimetic N-*tert*-butylhydroxylamine (NtBuHA) attenuated autophagy inhibition associated with GNS561 (Fig. 5C), indicating that inhibition of PPT1 function by GNS561 induced the observed anti-autophagy effect.

**Fig. 5.**
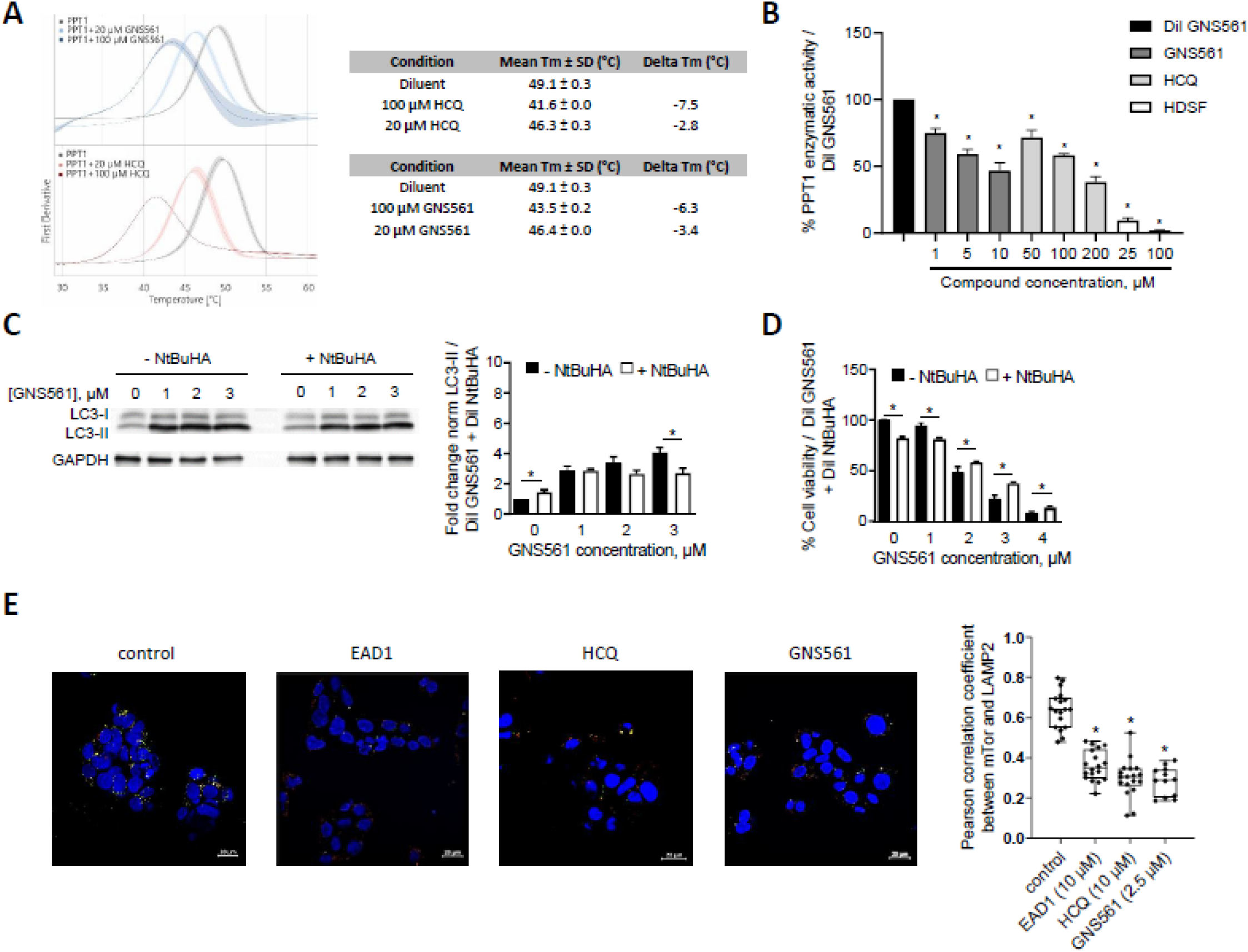
GNS561 targets palmitoyl-protein thioesterase 1 (PPT1). (A) Nano differential scanning fluorimetry assays comparing GNS561 + PPT1 and hydroxychloroquine (HCQ) + PPT1 against the apo-PPT1 ligand. Data represent the mean (solid lines) ± SEM (shaded areas) of two experiments. (B) PPT1 enzymatic activity of HepG2 cells treated with GNS561 for 3 h. HCQ and hexadecanesulfonyl fluoride (HDSF) were used as positive controls. The results were compared to the diluent of GNS561 (control condition). (C) Representative immunoblotting of light chain 3 phosphatidylethanolamine conjugate (LC3-II) in HepG2 cells treated with GNS561 for 16 h in the presence or absence of NtBuHA (8 mM). Glyceraldehyde-3-phosphate dehydrogenase (GAPDH) was used as a loading control. Fold changes of normalized LC3-II level were calculated against the control condition (diluent of GNS561 + diluent of NtBuHA). (D) Cell viability percent against the control condition (diluent of GNS561 + diluent of NtBuHA) after 24 h of treatment with GNS561 in the presence or absence of NtBuHA (8 mM). (E) Staining of lysosomes (lysosomal-associated membrane protein 2 [LAMP2], green), mTOR (red) and nucleus (4′,6-diamidino-2-phenylindole [DAPI], blue) after treatment with GNS561 and two positive controls, EAD1 and HCQ, for 16 h. Pearson correlation coefficient between mTOR and LAMP2 was represented using box and whisker representation (min to max). Scale bars represent 20 μm. In (B), (C) and (D), data represent the mean + SEM. For comparison, Student t-test was used for (C) and (D) and one-way ANOVA with Dunnett’s post hoc analysis was performed for (B) and (E). For all studies except (A), n ≥ 3 biological replicates. * represents significant difference, at least p < 0.05.

To determine whether inhibition of PPT1 function was responsible for the antitumoral activity of GNS561, HepG2 cells were treated with GNS561 with or without NtBuHA pretreatment. As shown in Fig. 5D, NtBuHA partially prevented the antitumor activity of GNS561, as evidenced by the increased viability of cells pretreated with NtBuHA. The same rescue effect of NtBuHA pretreatment was observed for HCQ used as a positive control (Supplementary Fig. 6). After demonstrating that NtBuHA had no impact on GNS561 lysosomal localization (Supplementary Fig. 7), we validated that the impact of NtBuHA pretreatment was due to its PPT1 mimetism, suggesting that inhibition of PPT1 function by GNS561 was partially liable for its antitumoral activity. The results of Rebecca et al. suggested that PPT1 inhibition could result in mTOR inhibition through the displacement of mTOR from the lysosomal membrane (4, 5). Thus, we investigated the localization of mTOR after GNS561 treatment using immunofluorescence microscopy. HCQ and EAD1 were used as positive controls (15). As shown in Fig. 5E, GNS561 treatment, as well HCQ and EAD1 treatments, significantly impaired mTOR localization to the lysosomal surface. Therefore, GNS561-induced PPT1 inhibition resulted in displacement of mTOR from the lysosomal membrane and consequently likely inhibited the mTOR signaling pathway.

### GNS561 induces LMP and cathepsin-dependent cell death

To characterize GNS561-induced changes in lysosomes, we analyzed LMP. To this end, we took advantage of the steady endocytic capacity of cells to load fluorescent dextran into lysosomes and the translocation of lysosomal localized dextran into the cytosol after LMP-inducing insult. Fluorescent dextran in healthy cells appears in dense punctate structures representing intact lysosomes, whereas after LMP, a diffuse staining pattern throughout the cytoplasm is seen. After GNS561 treatment, such diffuse dextran staining was observed (Fig. 6A), suggesting an induction of LMP. As seen in Fig. 6B, the loss of membrane integrity, which is the hallmark of LMP, was observed by transmission electron microscopy of HepG2 cells treated with 3 μM GNS561 for 24 h. To confirm this effect, cathepsin localization was studied after GNS561 treatment. After 48 h of treatment, GNS561 decreased cathepsin staining (Fig. 6C), indicating that cathepsins were released into the cytosol, thus validating LMP.

**Fig. 6.**
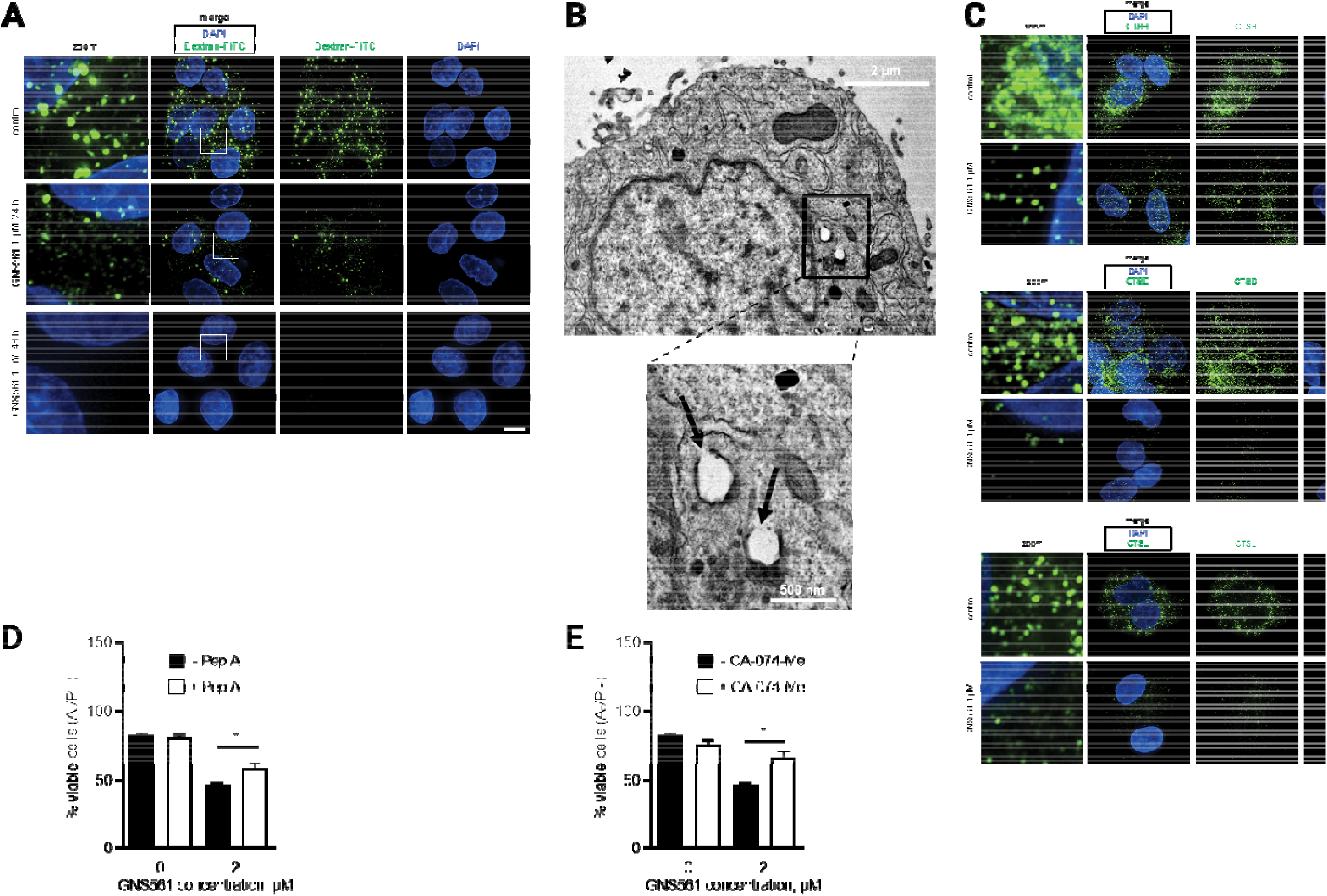
GNS561 induces lysosomal membrane permeabilisation and cathepsin-dependent cell death in HepG2 cells. (A) Localization of FITC-dextran after GNS561 treatment for the indicated times. (B) Electron microscopy imaging of lysosomal membrane permeabilization (arrows) after GNS561 treatment (3 μM) for 24 h. (C) Localization of cathepsin B (CTSB), cathepsin D (CTSD) and cathepsin L (CTSL) after GNS561 treatment for 48 h. (D) Viable cell (A-/PI-) analysis by flow cytometry of cells pretreated or not with pepstatin A (Pep A) (5 μM) for 1 h and then treated with Pep A (5 μM) and GNS561 or with GNS561 alone for 48 h. (E) Viable cell (A-/PI-) analysis by flow cytometry of cells pretreated or not with CA-074-Me (20 μM) for 1 h and then treated with CA-074-Me (20 μM) and GNS561 or with GNS561 alone for 48 h. Scale bars in (A) and (C) represent 10 μm. For all studies, n ≥ 3 biological replicates. Data represent the mean + SEM. For comparison, Student t-test was used. * represents significant difference, at least p < 0.05.

As cathepsin release into the cytosol after LMP may trigger cytosolic cellular death signaling (16), we evaluated the role of cathepsins in GNS56-induced cell death. To this end, HepG2 cells were pretreated with an inhibitor of CTSD, pepstatin A, or an inhibitor of CTSB, CA-074-Me. Under these conditions, cell viability was partially rescued (Fig. 6D and E), suggesting that the GNS561-induced apoptotic pathway is at least partially cathepsin-dependent.

## Discussion

Rapidly dividing and invasive cancer cells are strongly dependent on effective lysosomal functions. Lysosomes are acidic and catabolic organelles that are responsible for the disposal and recycling of used and damaged macromolecules and organelles, as well as the assimilation of extracellular materials incorporated into the cell by endocytosis, autophagy, and phagocytosis. Increased autophagic flux and changes in lysosomal compartments in cancer cells have been shown to promote invasion, proliferation, tumor growth, angiogenesis, and drug resistance. Consistently, lysosomal changes are expected to sensitize cells to lysosome-targeting anticancer drugs (17). Many steps in the autophagy pathway represent potentially druggable targets and several clinical trials have aimed to block autophagy by inhibiting lysosomal functions using CQ and HCQ. Unfortunately, CQ and HCQ failed to demonstrate consistent antitumor effects possibly due to subeffective anticancer concentrations in humans, even with high doses. Drug screening led us to identify GNS561 as a lead compound that displays lysosomotropism and significantly higher antiproliferative effects in human cancer cells compared to HCQ.

We previously reported that GNS561 yielded antiproliferative activity in iCCA, inhibited late-stage autophagy, and induced a dose-dependent enlargement of lysosomes (11). Based on these preliminary results, we further investigated the cellular mechanisms by which GNS561 may lead to lysosomal changes and death in cancer cells. In this study, we confirmed that GNS561 antitumor properties are strongly dependent on its lysosomotropic properties. In accordance with the hypothesis proposed in our previous study (11), we showed here that GNS561 induced a dose-dependent increase in the number of enlarged lysosomes and LMP leading to cytosolic cathepsin release, caspase activation, and apoptotic cell death. These observations confirm prior reports that highlight the capability of lysosomotropic agents to cause lysosomal stress and lysosomal enlargement (18). Further investigations are needed to identify the upstream signals that initiate LMP in GNS561-treated cells.

PPT1 plays a central role in the control of cellular autophagy by enabling the degradation and intracellular trafficking of membrane-bound proteins. PPT1 is highly expressed in several cancer cell lines as well as in advanced stage cancers in patients (4). Recent data have shown that lysosome-specific inhibitors targeting PPT1 can modulate protein palmitoylation and display antitumor activity in melanoma and colon cancer models (5). Our data showed that PPT1 acts as a molecular target of GNS561. GNS561 bound to PPT1 and inhibited its activity in cells. Cells treated with the chemical mimetic NtBuHA were partially resistant to GNS561-mediated anticancer effect and attenuated GNS561-associated autophagic flux inhibition, suggesting that inhibition of the thioesterase activity of PPT1 is essential for the anti-autophagic and antitumoral effects of GNS561. It is to highlight that as showed in studies of Amaravadi team, a partial inhibition of PPT1 activity (25% at 1 μM GNS561, 3 h) finally resulted in strong decrease of cell viability (more than 50% at 1 μM GNS561, 72 h).

Moreover, we observed that GNS561 modified the intracellular localization of mTOR. This is in accordance with previous studies showing that inhibition of PPT1 may displace the mTOR protein from the lysosomal membrane as a result of the inhibition of vATPase/Ragulator/Rag GTPase interactions (4, 5). It was also described that lysosomal mTOR localization brings it in close vicinity to its main regulator, Rheb, and that as a result, the mTOR/Rheb interaction can activate mTOR kinase activity leading to the phosphorylation of downstream effectors (19). Consistently, we hypothesized that GNS561-induced PPT1 inhibition led to mTOR signaling pathway inhibition. Further investigations on downstream targets of mTOR as S6 and 4E-BP1 kinases are currently ongoing to fully validate this hypothesis.

As previously observed in iCCA (11), we showed here that GNS561 induced a significant decrease in the enzymatic activity of cathepsins. This decreased activity is unlikely due to a direct inhibition of CTSL, CTSB and CTSD by GNS561 but rather could be the consequence of both impairment of CTSL and CTSD maturation and lysosomal unbound Zn^2+^ accumulation. As cathepsin activity is optimal in acidic pH (20), we could also speculate that GNS561 may negatively influence the proteolytic activity of cathepsins by inducing an increase in lysosomal pH via PPT1 inhibition. In fact, other authors have shown that PPT1 deficiency in Cln1−/− mice disrupted the delivery of the v-ATPase subunit V0a1 to the lysosomal membrane, leading to a dysregulation of lysosomal acidification (21). The authors suggested that S-palmitoylation by PPT1 may play a critical role in the trafficking of the V0a1 subunit of v-ATPase to the lysosomal membrane and in lysosomal pH regulation.

Based on prior studies, GNS561 was neither a zinc ionophore nor a zinc chelator (data not shown), unlike CQ (22). However, our hypothesis that GNS561-induced PPT1 inhibition could lead to lysosomal deacidification could also explain the observed lysosomal unbound Zn^2+^ accumulation after GNS561 treatment. In fact, as lysosomal pH is mainly regulated by cation/anion movement across the lysosomal membrane, it was suggested that a proton motive force was required to mediate unbound Zn^2+^ efflux (12).

In summary, GNS561-induced PPT1 inhibition may lead to two main mechanisms inducing cancer cell death. One is related to lysosomal deacidification, which induces lysosomal unbound Zn^2+^ accumulation, a decrease in the enzymatic activity of cathepsins, inhibition of autophagic flux, lysosomal swelling, LMP, cathepsin release, and caspase-dependent apoptosis. The other is linked to prevention of the interaction between v-ATPase and the Ragulator complex, blockage of mTOR lysosomal recruitment, impairment of mTOR–Rheb interaction and finally the inhibition of mTOR signaling pathway. Thus, by targeting PPT1, GNS561 acts as a regulator of autophagy and mTOR, two major processes that drive cancer aggressiveness. Finally, as lysosomes and autophagy are associated with adaptive mechanisms of resistance to mTOR inhibition (23), GNS561 can disable mTOR function and downregulate adaptive mechanisms of resistance.

An extensive preclinical program has been conducted to evaluate the antitumor activity, pharmacological properties and toxicology of GNS561. Our data showed that GNS561 displays antiproliferative effects in several human cancer cells (cell lines and primary patient-derived cells) and that GNS561 was more potent than HCQ. Analysis of the whole-body tissue distribution of GNS561 in rats after repeated oral dosing of GNS561 showed that GNS561 was mainly concentrated in the liver, stomach and lung. The data are consistent with the basic lipophilic nature of GNS561 and with studies showing that basic lipophilic drugs show high lysosomal tropism and high uptake in lysosome-profuse tissues, such as the liver and the lung (24). As GNS561 had a high liver tropism, the effect of GNS561 on tumor growth in vivo was evaluated using two liver cancer models: one orthotopic human liver cancer xenograft mouse model (with an HCC patient-derived cell line, LI0752) and one DEN-induced cirrhotic rat model with HCC. These studies showed that GNS561 administered by oral gavage was well tolerated up to the doses of 50 mg/kg/day for 6 days in mice and up to 15 mg/kg/day for 6 weeks in rats and induced significant antitumor growth activity that was either comparable to or higher than sorafenib. In addition, in a DEN-induced cirrhotic rat model with HCC, the combination of GNS561 with sorafenib exerted an additive effect in controlling tumor progression and cell proliferation. This result was in accordance with a study completed by Shimizu et al. showing that sorafenib increased autophagy in some HCC cells, which leads to resistance and that combination of sorafenib with an autophagy inhibitor significantly increase the suppression of tumor growth in vivo (25). Further studies are required to investigate GN561 mechanism of action in vivo and particularly to confirm the engagement of PPT1 in GNS561 antitumor efficacy in vivo.

Furthermore, instead of that observed with CQ and HCQ (26), the distribution of GNS561 into the central nervous system was limited. Inactivating PPT1 mutations have long been known to induce infantile neuronal cerebral lipofuscinosis and induce retinopathy during childhood (27). Germline PPT1 mutations were shown to selectively affect the central nervous system, with no effects in other tissues. Prior clinical experience using CQ and HCQ showed that retinopathy was one of the major toxicities in patients (28). Authors have suggested that novel PPT1 inhibitors may take advantage of not crossing the blood-brain barrier to avoid retinal toxicity (4). Interestingly, our data shown that the disposition of GNS561 displays limited penetration into the brain in rats, consistent with the lack of neurological and retinal toxicity observed in the current Phase 1b/2a clinical trial of GNS561 (29, 30). In brief, our findings strengthen the importance of PPT1 and lysosomes as cancer targets. Recently, it was shown that PPT1 inhibition by CQ derivatives or genetic Ppt1 inhibition increases the antitumor activity of anti-PD-1 antibody in melanoma by M2 to M1 phenotype switching in macrophages and a reduction in myeloid-derived suppressor cells in the tumor (31). As such, GNS561 represents a promising new candidate for drug development in HCC either alone or in combination with other drugs, such as anti-PD-1 antibody.

## List of abbreviations

CTS: cathepsin
CQ: chloroquine
iCCA: intrahepatic cholangiocarcinoma
DEN: diethylnitrosanime
GAPDH: glyceraldehyde-3-phosphate dehydrogenase
HCC: hepatocellular carcinoma
HCQ: hydroxychloroquine
LMP: lysosomal membrane permeabilization
NtBuHA: N-tert-butylhydroxylamine
PPT1: palmitoyl-protein thioesterase 1

## Acknowledgments

The authors are very grateful to Dr. Sebastian Müller and Dr. Raphaël Rodriguez from Curie Institute for mechanistic analysis, Pr. Thierry Levade and Dr. Nathalie Andrieu from CRCT for the PPT1 enzymatic assay, Keerthi Kurma and Seyedeh Tayebeh Ahmad Pour from the Institute for Advanced Biosciences for technical support during animal experiments and Dr. François Autelitano, Dr. Marie Guillemot and Philippe Fabre for Zn^2+^ localization analysis.

## References

1. Bray F, Ferlay J, Soerjomataram I, Siegel RL, Torre LA, Jemal A. Global cancer statistics 2018: GLOBOCAN estimates of incidence and mortality worldwide for 36 cancers in 185 countries. CA: a cancer journal for clinicians 2018;68:394–424.

2. Aits S, Jaattela M. Lysosomal cell death at a glance. Journal of Cell Science 2013;126:1905–1912.

3. Klempner SJ, Myers AP, Cantley LC. What a tangled web we weave: emerging resistance mechanisms to inhibition of the phosphoinositide 3-kinase pathway. Cancer Discovery 2013;3:1345–1354.

4. Rebecca VW, Nicastri MC, Fennelly C, Chude CI, Barber-Rotenberg JS, Ronghe A, et al. PPT1 Promotes Tumor Growth and Is the Molecular Target of Chloroquine Derivatives in Cancer. Cancer Discovery 2019;9:220–229.

5. Rebecca VW, Nicastri MC, McLaughlin N, Fennelly C, McAfee Q, Ronghe A, et al. A Unified Approach to Targeting the Lysosome’s Degradative and Growth Signaling Roles. Cancer Discovery 2017;7:1266–1283.

6. Shi T-T, Yu X-X, Yan L-J, Xiao H-T. Research progress of hydroxychloroquine and autophagy inhibitors on cancer. Cancer Chemotherapy and Pharmacology 2017;79:287–294.

7. Verbaanderd C, Maes H, Schaaf MB, Sukhatme VP, Pantziarka P, Sukhatme V, et al. Repurposing Drugs in Oncology (ReDO)-chloroquine and hydroxychloroquine as anti-cancer agents. Ecancermedicalscience 2017;11:781.

8. Manic G, Obrist F, Kroemer G, Vitale I, Galluzzi L. Chloroquine and hydroxychloroquine for cancer therapy. Molecular & Cellular Oncology 2014;1:e29911.

9. Pérez-Hernández M, Arias A, Martínez-García D, Pérez-Tomás R, Quesada R, Soto-Cerrato V. Targeting Autophagy for Cancer Treatment and Tumor Chemosensitization. Cancers 2019;11:1599.

10. Plantone D, Koudriavtseva T. Current and Future Use of Chloroquine and Hydroxychloroquine in Infectious, Immune, Neoplastic, and Neurological Diseases: A Mini-Review. Clinical Drug Investigation 2018;38:653–671.

11. Brun S, Bassissi F, Serdjebi C, Novello M, Tracz J, Autelitano F, et al. GNS561, a new lysosomotropic small molecule, for the treatment of intrahepatic cholangiocarcinoma. Investigational New Drugs 2019;37:1135–1145.

12. Lockwood TD. Lysosomal metal, redox and proton cycles influencing the CysHis cathepsin reaction. Metallomics: Integrated Biometal Science 2013;5:110–124.

13. Lockwood TD. Biguanide is a modifiable pharmacophore for recruitment of endogenous Zn2+ to inhibit cysteinyl cathepsins: review and implications. Biometals: An International Journal on the Role of Metal Ions in Biology, Biochemistry, and Medicine 2019;32:575–593.

14. Turk V, Stoka V, Vasiljeva O, Renko M, Sun T, Turk B, et al. Cysteine cathepsins: From structure, function and regulation to new frontiers. Biochimica et Biophysica Acta (BBA) - Proteins and Proteomics 2012;1824:68–88.

15. Sironi J, Aranda E, Nordstrøm LU, Schwartz EL. Lysosome Membrane Permeabilization and Disruption of the Molecular Target of Rapamycin (mTOR)-Lysosome Interaction Are Associated with the Inhibition of Lung Cancer Cell Proliferation by a Chloroquinoline Analog. Molecular Pharmacology 2019;95:127–138.

16. Oberle C, Huai J, Reinheckel T, Tacke M, Rassner M, Ekert PG, et al. Lysosomal membrane permeabilization and cathepsin release is a Bax/Bak-dependent, amplifying event of apoptosis in fibroblasts and monocytes. Cell Death and Differentiation 2010;17:1167–1178.

17. Kallunki T, Olsen OD, Jäättelä M. Cancer-associated lysosomal changes: friends or foes? Oncogene 2013;32:1995–2004.

18. Wang F, Gómez-Sintes R, Boya P. Lysosomal membrane permeabilization and cell death. Traffic 2018;19:918–931.

19. Carroll B, Maetzel D, Maddocks OD, Otten G, Ratcliff M, Smith GR, et al. Control of TSC2-Rheb signaling axis by arginine regulates mTORC1 activity. eLife 2016;5.

20. Turk B, Dolenc I, Lenarcic B, Krizaj I, Turk V, Bieth JG, et al. Acidic pH as a physiological regulator of human cathepsin L activity. European Journal of Biochemistry 1999;259:926–932.

21. Bagh MB, Peng S, Chandra G, Zhang Z, Mukherjee AB. Dysregulation of V-ATPase Function Impairs Lysosomal Acidification in a Mouse Model of a Rare Lysosomal Storage Disease, INCL. The FASEB Journal 2016;30:879.871–879.871.

22. Xue J, Moyer A, Peng B, Wu J, Hannafon BN, Ding W-Q. Chloroquine Is a Zinc Ionophore. PLoS ONE 2014;9:e109180.

23. Xie X, White EP, Mehnert JM. Coordinate Autophagy and mTOR Pathway Inhibition Enhances Cell Death in Melanoma. PLOS ONE 2013;8:e55096.

24. Daniel WA, Wójcikowski J. Contribution of lysosomal trapping to the total tissue uptake of psychotropic drugs. Pharmacology & Toxicology 1997;80:62–68.

25. Shimizu S, Takehara T, Hikita H, Kodama T, Tsunematsu H, Miyagi T, et al. Inhibition of autophagy potentiates the antitumor effect of the multikinase inhibitor sorafenib in hepatocellular carcinoma. Int J Cancer 2012;131:548–557.

26. Harder BG, Blomquist MR, Wang J, Kim AJ, Woodworth GF, Winkles JA, et al. Developments in Blood-Brain Barrier Penetrance and Drug Repurposing for Improved Treatment of Glioblastoma. Frontiers in Oncology 2018;8.

27. Metelitsina TI, Waggoner DJ, Grassi MA. BATTEN DISEASE CAUSED BY A NOVEL MUTATION IN THE PPT1 GENE:. Retinal Cases & Brief Reports 2016;10:211–213.

28. Marmor MF, Kellner U, Lai TYY, Lyons JS, Mieler WF, Ophthalmology AAo. Revised recommendations on screening for chloroquine and hydroxychloroquine retinopathy. Ophthalmology 2011;118:415–422.

29. Eide DJ. The SLC39 family of metal ion transporters. Pflugers Arch 2004;447:796–800.

30. ClinicalTrials.gov. Study of GNS561 in Patients With Liver Cancer. In: ClinicalTrials.gov.

31. Sharma G, Ojha R, Noguera-Ortega E, Rebecca VW, Attanasio J, Liu S, et al. PPT1 inhibition enhances the antitumor activity of anti-PD-1 antibody in melanoma. JCI insight 2020;5.

