## Supplementary data for "GNS561, a clinical-stage PPT1 inhibitor, is efficient against hepatocellular carcinoma via modulation of lysosomal functions"

### Supplementary Materials and Methods

#### *Reagents and antibodies*

Bafilomycin A1 (BafA1) (#254134-5G), ammonium chloride ($NH_4Cl$) (#254134-5G), E-64 (L-3-carboxy-trans-2,3-epoxy-propionyl-leucylamide-(4-guanido)-butane) (#3132), CA-074 (N-(L-3-trans-propylcarbamoyloxirane-2-carbonyl)-L-isoleucyl-L-proline) (#C5732-5MG), Dextran-FITC (#FD10S), Pepstatin A (#P5318-25MG), dithiothreitol (#D0632), diethylnitrosamine (DEN) (#N0756), human cathepsin D (CTSD) (#C8696-25UG), almond β-glucosidase (#G4511), 4-methylumbelliferone (#M1381), N-(tert-Butyl)hydroxylamine acetate (NtBuHA) (#479675), CA-074-Me (C5732-5MG) and hydroxychloroquine sulfate (HCQ) (#H0915) were obtained from Sigma-Aldrich (MO, USA). Human cathepsin L (CTSL) (#219402-25UG), human cathepsin B (CTSB) (#219364-5U) and the fluorogenic substrates Z-Phe-Arg-7-amido-4-methylcoumarin (#03-32-1501), Z-Arg-Arg-7-amido-4-methylcoumarin (#219392) and (7-methoxycoumarin-4-yl)acetyl-Gly-Lys-Pro-Ile-Leu-Phe-Phe-Arg-Leu-Lys(Dnp)-D-Arg-$NH_2$ (#219360) were supplied by Merck (Germany). Mammalian Cell Lysis Buffer (GE Healthcare, Chicago, IL, USA, #28-9412-79), 4-methylumbelliferyl-β-D-6-thiopalmitoyl-glucoside (Moscerdam, Netherlands, #EM06650), Z-VAD-FMK (Bio Techne, : Minneapolis, MN, USA, #FMK001), sorafenib (Santa Cruz Biotechnology, Dallas, TX, USA, #Sc-357801), hexadecanesulfonyl fluoride (HDSF) (Santa Cruz Biotechnology, #SC-221708), human PPT1 (OriGene Technologies, Rockville, Md., USA, #TP721098), DC661 (Vagdevi Innoscience, India), Triton X-100 (Dutscher, France, #091584B) and cOmplete™ Protease Inhibitor Cocktail (Roche, Switzerland, #4693132001) were used. Fluozin-3 (#F24195), LysoTracker Deep Red (#L12492), CaspGLOW™

Fluorescein Active Staining Kit (#88-7003-42), Annexin V/PI kit (#V13242) were purchased from Thermo Fisher Scientific (Waltham, MA, USA).

For immunoblotting assays, rabbit anti-light chain 3 phosphatidylethanolamine conjugate (LC3-II) (Sigma-Aldrich, # L7543, 1:3,000), mouse anti-glyceraldehyde-3-phosphate dehydrogenase (GAPDH) (Abnova, Taïwan, #H00002597-M01, 1:5,000), rabbit anti-Poly(ADP-ribose) polymerase (PARP) (GeneTex, Irvine, CA, USA, #GTX100573, 1:1,500), rabbit anti-cleaved caspase-3 (Asp175) (Cell Signaling, Danvers, MA, USA, #9661, 1:200), goat anti-CTSD (Santa Cruz Biotechnology, #Sc-6486, 1:200), rabbit anti-CTSB (Santa Cruz Biotechnology, #Sc-13985, 1:200), goat anti-CTSL (Santa Cruz Biotechnology, #Sc-6498, 1:200), goat anti-rabbit (Santa Cruz Biotechnology, #Sc-2004, 1:5,000), donkey anti-goat (Santa Cruz Biotechnology, #Sc-2020, 1:5,000), goat anti-mouse (Jackson ImmunoResearch, United Kingdom, #115-035-003, 1:40,000), goat anti-rabbit (Sigma, #AP307P, 1:25,000) and goat anti-rabbit (Jackson ImmunoResearch, #111-035-003, 1:40,000) antibodies were used.

For immunofluorescence assays, anti-LAMP1 (Cell Signaling, #9091S, 1:200), anti-LAMP2 (Developmental Studies Hybridoma Bank, Iowa City, IA, USA, #H4B4-S, 1:100), anti-mTor (Cell Signaling, #2972, 1:250), anti-CTSB (Abcam, #ab58802, 1:200), anti-CTSD (Abcam, #ab75852, 1:200), anti-CTSL (Abcam, #ab133641,1:200), Alexa Fluor 594 conjugate (Life Technologies, Carlsbad, CA, USA, #A-11072, #A-11032, 1:500), Alexa Fluor 488 conjugate (Life Technologies, #A-11017, #A-11008, 1:500), Alexa Fluor 647 conjugate (Life Technologies, #A-21447, 1:500) and Alexa Fluor 546 conjugate (Life Technologies, #A-11003, 1:1,000) antibodies were used. Antibodies were diluted in blocking solution (5% bovine serum albumin (BSA), 0.1% Tween-20/TBS or 5% fetal bovine serum (FBS) in PBS).

For immunohistochemical analysis, anti-Ki67 (Thermo Fisher Scientific, #MA5-14520, 1:150) and anti-Cyclin D1 (Abcam, #ab134175, 1:200) antibodies were used.

*Cell lines and cell culture*

Huh7 (hepatocellular carcinoma, JCRB Cell Bank, Japan, #JCRB0403), HCT-116 (colorectal carcinoma, ATCC, Manassas, VA, USA, #CCL-247), A549 (lung cancer, Sigma, #86012804), LN-18 (glioblastoma, ATCC, #CRL-2610), LN-229 (glioblastoma, ATCC, #CRL-2611), MDA-MB-231 (breast cancer, ATCC, #HTB-26) and A375 (malignant melanoma, ATCC, #CRL-1619) cell lines were cultured in Dulbecco’s modified Eagle’s medium (DMEM) (Dutscher, #L0103-500). KG-1 cell line (acute myeloid leukemia, ATCC, #CCL-246) was maintained in Iscove medium (Dutscher, #L0191-500). CAKI-1 (renal adenocarcinoma, NCI, New York, NY, USA, #0507829), 786-O (renal adenocarcinoma, NCI, #05007648), DU-145 (prostate cancer, ATCC, #HTB-81), PC-3 (prostate cancer, Sigma, #90112714), NCI-H358 (lung cancer, Sigma, #95111733) and NIH:OVCAR3 (ovarian adenocarcinoma, ATCC, #HTB-161) cell lines were maintained in RPMI 1640 medium (Dutscher, #L0498-500). HepG2 (hepatocellular carcinoma, ATCC, #HB-8065), HT-29 (colorectal carcinoma, ATCC, #HTB-38 ) and SK-MEL-28 (malignant melanoma, ATCC, #HTB-72) cell lines were cultured in Dulbecco’s modified Eagle’s medium low glucose (Thermo Fisher Scientific, #21885025), Mc Coy's medium (Dutscher, #L0210-500), DMEM: Nutrient Mixture F-12 (Thermo Fisher Scientific, #31765-027) and MEM (Dutscher, #L0416-500) respectively.

Cell lines were tested for Mycoplasma at each thawing and were used at a number of passage lower than 20. All cell lines were maintained in medium containing 1%

penicillin-streptomycin (Dutscher, #P06-07100) and 10% FBS (GE Healthcare, #SV30160.03C), except NIH:OVCAR3 and KG-1 cell lines that were cultured in medium supplemented 20% FBS. In NIH:OVCAR3 medium, 0.01 mg/mL insulin was also added.

*Cell viability assay*

Cell viability was assessed using the CellTiter-Glo Luminescent Cell Viability Assay following the manufacturer's protocol (Promega, Madison, WI, #G7573). Briefly, cells were plated in a 96-well tissue culture plate in 90 µL of medium. Twenty-four hours after plating, cells were treated with 10 µL of increasing concentrations of GNS561 or with GNS561 vehicle and were incubated for 72 h. At the end of the treatment, 100 µL of CellTiter-Glo solution was added to each well; cells were briefly shaken and then were incubated at room temperature (RT) for 10 min to allow stabilization of the luminescent signal. The luminescence was recorded using an Infinite F200 Pro plate reader (Tecan, Switzerland) and cell viability was expressed as a percentage of the values obtained from the negative control cells (vehicle treated cells). The half-maximal inhibitory concentration ($IC_{50}$) was evaluated using a nonlinear regression curve in Prism 8.4.3 software (GraphPad Software, La Jolla, CA, USA). Each concentration was tested in triplicate. Mean $IC_{50}$ was calculated as the average of three independent experiments.

*Flow cytometry for annexin V/*propidium *iodide assay*

Cells were treated as indicated in the figures. Cell media containing floating cells were recovered and pelleted by centrifugation at 300 × g. Cells were trypsinized and recovered in medium and pelleted by centrifugation at 300 × g. Both cell pellets were combined and washed twice with ice cold PBS. Annexin V and propidium iodide staining was performed according to the manufacturer’s protocol. Cells were analyzed immediately using flow cytometry and data were recorded on a BD Accuri C6 (BD Biosciences, San Jose, CA, USA) and processed using Cell Quest or BD FACSDiva™ softwares (BD Biosciences) or FlowJo software (FLOWJO, USA).

*Flow cytometry detection of caspase activity*

For these experiments the manufacturer’s protocol of CaspGLOW Fluorescein Active Staining Kit was followed. In brief, cells were collected, washed with PBS and stained with Z-VAD-FMK-FITC. Cells were kept on ice and analyzed immediately using flow cytometry and data were recorded on a BD Accuri C6 (BD Biosciences) and processed using BD Cell Quest (BD Biosciences) and FlowJo softwares.

*Luminescence detection of caspase activity*

The activity of caspase 3/7 and caspase 8 was measured using the Caspase-Glo 3/7 Assay (Promega, #G8092) and Caspase-Glo 8 Assay (Promega, #G8202) following the manufacturer’s protocol. Briefly, HepG2 cells were plated in a 96-well plate (7,500 cells per well) in 90 µL of medium. Twenty-four hours after plating, cells were treated with 10 µL of GNS561 (1-4 µM) or GNS561 vehicle and incubated for 6, 24 and 30 h. At the end of the treatment, 100 µL of Caspase-Glo 3/7 or Caspase-Glo 8 reagent were added to each well and cells were incubated for 1 h at RT. Then, luminescence was measured by an Infinite F200 Pro plate reader. Fold change of activation of caspase 3/7 and caspase 8 was determined by comparing the luminescence in the treated groups with the luminescence observed in the negative control wells (vehicle treated cells), with the luminescence of blank wells subtracted. At each time point, in parallel with the activation of caspase 3/7 and caspase 8, cell viability was also investigated using CellTiter-Glo Luminescent Cell Viability Assay. Each GNS561 concentration was tested in triplicate in three independent experiments.

*Z-VAD-FMK, CA-074-Me, Pepstatin A, NtBuHA, Baf A1, $NH_4Cl$ and GNS561 treatments*

On the day of the treatment, cells were treated with warmed media and specified concentrations of Z-VAD-FMK (5 µM, pretreatment for 1 h), CA-074-Me (20 µM, pretreatment for 1 h), Pepstatin A (5 µM, pretreatment for 1 h), NtBuHA (8 mM), BafA1 (200 nM, pretreatment for 2 h), $NH_4Cl$ (20 mM, pretreatment for 2 h) and GNS561 (concentrations and time as indicated on the figures) or vehicle were added. Cells were treated for the times indicated and processed for several analyses as indicated. Each condition was tested in triplicate, and three independent experiments were performed.

*Chemical labeling of GNS561D in cells*

Cells were cultured at ~80% confluence and were treated with 10 µM GNS561D, a photoactivable analogue of GNS561 (with a diazide moiety), for 90 min. Cell were fixed with formaldehyde (2% in PBS, 12 min) prior to permeabilization (Triton X-100, 0.1% in PBS, 5 min) and washed three times with 1% BSA/PBS. The click reaction cocktail was prepared from Click-iT EdU Imaging kits (Life Technologies, #C10337) according to the manufacturer's protocol. Briefly, mixing 430 µL of 1× Click-iT reaction buffer with 20 µL of $CuSO_4$ solution, 1.2 µL Alexa Fluor azide, 50 µL reaction buffer additive (sodium ascorbate) to reach a final volume of 500 µL. Cover-slips were incubated with the click reaction cocktail in the dark at RT for 30 min, then washed three times with PBS. Immunofluorescence was then performed as indicated.

*Lysosome staining using Lysotracker*

LysoTracker Deep Red was added to the cells at the same time as GNS561 at 1:10,000.

*Cathepsin activity assay in cell lysate*

Twenty-four hours after HepG2 cell plating, the cells were treated with GNS561 (1, 2 and 4 µM) for 6 and 24 h. Treatment with vehicle was used as a baseline for cathepsin activity control. Cell lysates (1 µg of total protein) were pre-incubated with acetate buffer (0.1 M sodium acetate, pH 5.5, 10 mM DTT, 2 mM EDTA, and 0.01% Brij35) or with citrate buffer (0.1 M sodium citrate, pH 4.0, 2 mM EDTA, 0.01% Brij35) prior to measurement of the respective CTSB/L (including both cathepsins B and L) and CTSB activities and CTSD activity as previously reported (1). The peptidase activity of CTSB, CTSB/L and CTSD were determined fluorometrically with a fluorescence reader (Gemini spectrofluorometer, Molecular Devices, San José, CA, USA) using respectively, the synthetic substrates Z-Arg-Arg-7-amido-4-methylcoumarin (excitation wavelength: 350 nm; emission wavelength: 460 nm), Z-Phe-Arg-7-amido-4-methylcoumarin (excitation wavelength: 350 nm; emission wavelength: 460 nm) and 7-methoxycoumarin-4-yl)acetyl-Gly-Lys-Pro-Ile-Leu-Phe-Phe-Arg-Leu-Lys(Dnp)-D-Arg-$NH_2$ (excitation wavelength: 325 nm; emission wavelength: 390 nm) (2). The synthetic protease inhibitors E-64 (pan-inhibitor of cysteine cathepsins), CA-074 (specific inhibitor of CTSB) and Pepstatin A (inhibitor of CTSD) were used as controls to confirm the detection of specific activities. Slopes of the enzymatic activities were calculated with the software SoftMax Pro (Molecular Devices). For each experiment and tested condition, fold change of the cathepsin activity was determined by comparing the slope of the enzymatic activity in treated conditions to the slope of the enzymatic activity in the vehicle condition. Three independent experiments were performed.

*Unbound zinc staining using Fluozin*

Fluozin-3 was added for 30 min to live cells at 1:1,000.

*Activity of recombinant cathepsins B, L and D*

Recombinant CTSB (0.5 nM) was incubated at 30 °C in 0.1 M sodium acetate buffer, pH 5.5, containing 2 mM EDTA, 10 mM DTT and 0.01% Brij35, in the presence of GNS561 (0, 4 and 10 µM) for 30 min, before measurement of the residual enzymatic activity using 20 µM Z-Phe-Arg-7-amido-4-methylcoumarin as substrate. The same protocol was repeated for recombinant CTSL (2 nM). Alternatively, recombinant CTSD (0.5 nM) was incubated at 30 °C in 0.1 M sodium citrate buffer, pH 4.0, EDTA 2mM, Brij 35 0.01% in the presence of GNS561 (0, 4 and 10 µM) for 30 min, before measurement of the residual peptidase activity using 20 µM 7-methoxycoumarin-4-yl)acetyl-Gly-Lys-Pro-Ile-Leu-Phe-Phe-Arg-Leu-Lys(Dnp)-D-Arg-$NH_2$ as substrate (3). The synthetic protease inhibitors E-64 (pan-inhibitor of cysteine cathepsins) and Pepstatin A (inhibitor of CTSD) were used as controls to confirm the detection of specific activities. Slopes were calculated and average values determined (software Softmaxpro, Molecular Devices). Assays were performed in triplicate.

*Western blotting*

In brief, cells were lysed with Mammalian Cell Lysis Buffer. cOmplete™ Protease Inhibitor Cocktail was added extemporaneously to the lysis buffer. Ten to twenty micrograms of protein from each cell lysate was separated on a 15% or 4-15% SDS-PAGE gel, transferred to a PVDF membrane, and blotted with antibodies. For all blots, GAPDH immunoblotting was used as a loading control. All the experiments were repeated at least three times. Representative autoradiograms are shown.

*Autophagy assay*

The autophagy pathway was studied as performed previously (4). Twenty-four hours after HepG2 or Huh7 cell plating, the cells were treated with GNS561 (0.5, 1 and 2 µM for HepG2 cell line and 0.4, 0.8 and 1.6 µM for Huh7 cell line) for 24 h. Treatment with vehicle was used as a baseline for autophagic flux control. In specified conditions, BafA1 (100 nM) was added for the last 2 h of treatment. The autophagic flux was calculated as the ratio between the LC3-II level normalized against GAPDH level (Norm LC3-II) with BafA1 and without BafA1.

*Nano differential scanning fluorimetry measurements*

Nano differential scanning fluorimetry was used to measure thermal stability of PPT1 in absence (0 µM ligand) and presence of the ligands, GNS561 and HCQ (used at 20 μM and 100 μM). Recombinant PPT1 was used at 3.6 μM in 1x PBS pH 7.4, 0.05% Tween-20. In order to facilitate comparability, the assay buffer was supplied with 1% DMSO in experiments using the DMSO-solved ligand GNS561. For each condition, a duplicate experiment was prepared and measured in standard capillaries (NanoTemper Technologies GmbH, Germany, #PR-C002) at 40% sensitivity, temperature range from 20-95°C, and a heating speed of 1°C/min on a Prometheus NT.48 instrument (NanoTemper Technologies GmbH). PPT1 unfolding was measured by detecting the temperature-dependent change in tryptophan/tyrosine fluorescence at emission wavelengths of 350 nm and 330 nm, respectively. Melting temperatures were determined by detecting the maximum of the first derivative of the fluorescence ratios (F350/F330). For this, an 8th order polynomial fit was applied to the transition region (PR.StabilityAnalysis_x64_1.0.3.10009, NanoTemper Technologies). For determination of the influence of ligands on PPT1 stability, ΔTm values were determined. Initially the standard deviation of the Tm of each condition was calculated from two replicates (n = 2). The average Tm of PPT1 (in the respective buffer) was subtracted by the average Tm of the respective compound condition to finally obtain ΔTm values of each compound condition. Tm shifts > 6*standard deviation of PPT1 (0.3°C) were considered as significant.

*PPT1 enzyme assay*

HepG2 cell line was treated with GNS561 (1, 5 and 10 μM), HCQ (50, 100 and 200 μM) and HDSF (25 and 100 μM) for 3 h. The cells were lysed in 0.5% Triton X-100 with cOmplete™ Protease Inhibitor Cocktail. The cell lysates were used a source of PPT1 and PPT1 activity was assayed using 4-methylumbelliferyl-β-D-6-thiopalmitoyl-glucoside as reported (5). Reaction mixtures contained 5 μL of cell lysate + 5 μL

0.5% Triton X-100 with cOmplete™ Protease Inhibitor Cocktail + 20 µl of substrate preparation (0.5 mM substrate, 1.5 mM dithiothreitol, 0.1 U almond β-glucosidase (Sigma, #G4511), 0.2% Triton X-100 and McIlvain's phosphate citrate buffer (0.2M $Na_2HPO_4$, 0.1 M citric acid). After 1 h incubation at 37 °C, the reaction was stopped by adding 200 µl of glycine/NaOH 0.5 M (pH 10.5) buffer. The amount of the released fluorescent product 4-methylumbelliferone was determined by fluorometry at 358 and 448 nm for the excitation and emission wavelengths, respectively. CNL1 (infantile subtype of ceroid lipofuscinosis) fibroblasts which contain mutations in the PPT1 gene and normal fibroblasts were used as control. 4-methylumbelliferone diluted in glycine/NaOH 0.5 M (pH 10.5) buffer was used to do a standard curve and to calculate the enzymatic activity of PPT1.

*Lysosomal membrane permeabilization assay*

HepG2 cells were plated 24 h prior to the experiments and then treated as indicated. Then, cells were treated with Dextran-FITC at 1 mg/ mL for 1 h in cell medium. Cells where then fixed with formaldehyde (2% in PBS, 12 min) and analyzed by fluorescence microscopy.

*Cell imaging*

For immunofluorescence, HepG2 cells were blocked with 2% BSA or 10% FBS supplemented with 0.2% Tween-20/PBS (blocking buffer) for 20 min at RT. Cover-slips were incubated with 50 to 100 µL of diluted primary antibodies in blocking buffer 1 h at RT. Cover-slips were then washed three times with blocking buffer and incubated as described above with the appropriate secondary antibodies for 1 h.

Cover-slips were washed three times with PBS and mounted using Mowiol (Sigma Aldrich, #81381) or Vectashield Mounting Medium with DAPI (VECTOR Labs, Burlingame, CA, USA, #H-1200). Fluorescence images were acquired using a Deltavision real-time microscope (Applied Precision) with 60×/1.4NA and 100×/1.4NA objectives or using a LSM 800 Airyscan confocal microscope (Zeiss, Germany) with a 63X oil objective. A typical z-stack of a field contained cells in the range of 3-10. Pearson correlation coefficients were determined using the ImageJ plugin Coloc 2 for each individual cell. The background (no cell) was set to 0 and pearson correlation coefficients were calculated above that threshold. In immunofluorescence quantifications, one point represents one cell.

For electron microscopy, HepG2 cells were fixed at least for 1 h with glutaraldehyde 2.5% in 0.1 M sodium cacodylate buffer. For resin embedding, cells were washed three times with a mixture of 0.2 M saccharose/0.1 M sodium cacodylate. Cells were then post-fixed for 1 h with 1% $OsO_4$ diluted in 0.2 M Potassium hexa-cyanoferrate (III) / 0.1 M sodium cacodylate solution. After three 10 min washes with distilled water, the cells were gradually dehydrated with ethanol by successive 10 min baths in 30, 50, 70, 96, 100, and 100% ethanol. Substitution was achieved by successively placing the cells in 25, 50, and 75% Epon solutions for 15 min. HepG2 cells were placed for 1 h in 100% Epon solution and in fresh Epon 100% over-night under vacuum at RT. Polymerization occured with cells in 100% fresh Epon for 72 h at 60°C. All solutions used above were 0.2 µm filtered. Ultrathin 70 nm sections were cut using a UC7 ultramicrotome (Leica) and placed on HR25 300 Mesh Copper/Rhodium grids (TAAB). Sections were contrasted according to Reynolds (Reynolds 1963). Electron micrographs were obtained on a Morgagni 268D (Philips / FEI) transmission electron microscope operated at 80 keV TEM.

*Whole body rat distribution of GNS561 by mass spectrometry imaging*

Sprague Dawley male rats (N=2) have been dosed orally with GNS561 at 40 mg/kg/day for 28 days with a single administration and were sacrificed 7 hours after the last administration. One rat received water and used as a control animal. After the sacrifice, whole body rats were shaved and their legs and tails sawed off. They were individually embedded in CMC 3% and fast-frozen in a dry ice/hexane mix then stored at -80°C before sample preparation for matrix assisted laser desorption ionization (MALDI) analysis. For each animal, 20 µm tissue sections through the sagittal section plan were performed on tape (Leica, Germany, #14041739651) in a cryomacrotome cryostat (Leica, #CM3600) at -20°C. Tissue sections on tape were cryodesiccated 24 h then stored at -80°C until use. Low resolution optical images of each slide were acquired using a standard office type scanner (Hewlett-Packard, Palo Alto, CA, USA).

Prior to start the MALDI matrix deposit, sections on tape were stuck on double conductive tape (XYZ-Axis Electrically Conductive Tape 9713, 3M) on MALDI target (Bruker, Billerica, MA, USA). 2,5-Dihydroxybenzoic acid solution was prepared at 40 mg/mL in methanol/$H_2O$ + 0.2% trifluoroacetic acid (1:1 v:v) for matrix deposition in the positive ion mode study. TM sprayer (HTX Imaging, Chapel Hill, NC, USA) was used for spraying the MALDI matrices over the tissue sections. Deuterated GNS561 was added to the MALDI matrix at 3 µM and used as an internal standard.

Sections were analyzed by MALDI imaging. MALDI images were obtained using a 7T MALDI-FTICR (Solarix, Bruker) equipped with a SmartBeam II laser used with a repetition rate of 1000 Hz, in positive ion mode. Mass spectra were acquired with a full scan mode of acquisition within the *m/z* 100-1,000 range at 650 µm of spatial resolution. The mass spectrum obtained for each position of the images corresponds to the averaged mass spectra of 300 consecutive laser shots on the same location. Prior to each data acquisition, external calibration was performed using endogenous compounds well known and MALDI matrix ions. FTMS Control 2.0 and FlexImaging 4.1 software packages (Bruker Daltonics) were used to control the mass spectrometer and set imaging parameters. Multimaging™ 1.1 software (ImaBiotech, France) was used to create the molecular distributions of GNS561 normalized by the internal standard.

*Animal treatment*

The animals were checked daily for clinical signs, effects of tumor growth and any other abnormal effects. For experiments involving the mouse model (performed in CrownBio [United Kingdom] facilities), the protocol and any amendment(s) or procedures involving the care and use of animals were reviewed and approved by the Institutional Animal Care and Use Committee of CrownBio prior to experimentation, and during the study, the care and use of animals was conducted in accordance with the regulations of the Association for Assessment and Accreditation of Laboratory Animal Care. For the rat model, all animals received humane care in accordance with the Guidelines on the Humane Treatment of Laboratory Animals (Directive 2010/63/EU), and experiments were approved by the animal Ethics Committee: GIN Ethics Committee No.004.

*Human HCC orthotopic patient-derived LI0752 xenograft mouse model*

Thirty six 8 weeks old BALB/c nude male mice (Biomedical Research Institute of Nanjing University) were housed in the animal facility of Science and Technology Park (Taicang Jiangsu). Tumor fragments from stock mice inoculated with selected primary human liver cancer tissues (LI0752 fragment (R21P9)) were harvested and used for inoculation into the left liver lobe of mice. Seven days after inoculation, mice were randomized in 3 groups (n=12/group) and treated by i) GNS561 (30 mg/kg), ii) GNS561 (50 mg/kg) or iii) vehicle, during following 34 days. On days 7 (for randomization purpose), 21, 28 and 41 post-inoculation, blood sample for alpha fetoprotein quantification in serum was collected. At the end of the treatment period, animals were euthanized by sodium pentobarbital overdose followed by cervical dislocation and tumor in the liver measured in two dimension using a caliper and weighted. Alpha fetoprotein serum levels from mice experiments were measured by an ELISA kit (Zhengzhou Biocell, China).

*DEN-induced cirrhotic rat model of HCC*

Thirty 6-week-old Fischer 344 male rats (Charles River Laboratories, Wilmington, MA, USA) were housed in the animal facility of Plateforme de Haute Technologie Animale (Jean Roget, University of Grenoble-Alpes, France). Rats were kept in individually ventilated cage systems at constant temperature and humidity with 2-3 animals in each cage having free access to food (standard diet) and water during the entire study period. Rats were treated weekly with intra-peritoneal injections of 50 mg/kg of DEN, which were diluted in olive oil in order to obtain a fully developed HCC on a cirrhotic liver after 14 weeks (6). Rats were randomized in 4 different groups and treated during six weeks by i) sorafenib (n=8), ii) GNS561 (n=8), iii) combination of GNS561 and sorafenib (n=6) or iv) rested untreated (control, n=8), as specified in Supplementary Fig. S2. Treatments of GNS561 (15 mg/kg/day), sorafenib (10 mg/kg/day) and combination (GNS561+sorafenib) were administered by oral gavage for a period of six weeks. The nutritional state was monitored by daily weighing of rats and protein-rich nutrition was added to the standard food in every cage where a loss of weight was observed. The food intake per cage was monitored during the last 6 weeks of the experiment. Food was withheld for 3-4 h before the animals were sacrificed.

Imaging study was conducted on a 4.7 Tesla MR Imaging system (BioSpec 47/40 USR, Bruker). All rats were subjected to 3 magnetic resonance imaging (MRI) scans: MRI1 was performed before randomization, MRI2 was performed after 3 weeks of treatment and MRI3 after 6 weeks of treatment. MRI analysis was done by an investigator who was blinded of treatment allocation.

After the third MRI scan, all rats were euthanized with vena cava blood sampling for hematologic and biochemical analyses. Each liver was weighed, the diameter of the five largest tumors was measured and the number of tumors larger than 1 mm on the surface of the liver was counted, all in a blinded manner. Tumor proliferation was studied by using anti-Ki67 and anti-Cyclin D1 antibodies.

Serum and plasma were tested for liver and kidney safety markers (albumin, alkaline phosphatase, alanine aminotransferase, aspartate aminotransferase, prothrombin time, total bilirubin, cholesterol, gamma-glutamyl transpeptidase, glucose, creatinine) by Charles River Clinical pathology Services using Olympus and Stago instruments. Alpha fetoprotein serum levels were measured by a Rat alpha FP ELISA kit (Aviva Systems Biology, San Diego, CA, USA, #OKEH00252) in rat serum. Liver and plasma triglycerides were analysed using a Triglycerides kit (Erba Mannheim, Germany, #120211).

*Statistical analysis*

Statistical analyses were performed using Prism 8.4.3 software. For datasets with normal distribution, multiple comparisons were performed using one-way ANOVA with Dunnett's post hoc analysis. The parametric Student t-test was used to compare two groups of data with normal distribution. Data are presented as the mean values ± standard error mean (SEM) unless stated otherwise. Statistical significance was defined as a p-value $< 0.05$ and has been indicated by an asterisk in all figures.

**Supplementary Table 1. Clinical and biological analyses of diethylnitrosanime-induced cirrhotic rat model with HCC.**

Rats were randomized in 4 different groups and treated during six weeks by i) sorafenib (n=8), ii) GNS561 (n=8), iii) combination of GNS561 and sorafenib (n=6) or iv) rested untreated (control, n=8). AFP, alpha fetoprotein; AST, aspartate aminotransferase; ALT, alanine aminotransferase; ALP, alkaline phosphatase; GGT, gamma-glutamyl transpeptidase; PT, prothrombin time; TB, total bilirubin. Data represent the mean ± SD. Significant difference compared to control: *, p <0.05. Significant difference compared to sorafenib: #, P <0.05.

| | Control | sorafenib | GNS561 | GNS561 + sorafenib |
|---|---|---|---|---|
| **Body weight** | | | | |
| % of initial weight | 4.9 ± 2.0 | 10.0 ± 0.6 | 0.6 ± 1.7 | 2.0 ± 1.2 |
| **Liver** | | | | |
| Weight (g) | 13.0 ± 0.4 | 13.1 ± 0.3 | 12.7 ± 0.5 | 11.2 ± 0.9 |
| Weight (% of body weight) | 4.70 ± 0.27 | 4.6 ±0.2 | 4.5 ± 0.1 | 3.9 ± 0.2 |
| Tumor growth (mg/g) | 31.6 ± 3.9 | 27.4 ± 1.9 | 27.5 ± 2.3 | 25.6 ± 1.8 |
| **Blood** | | | | |
| Albumin (g/dL) | 3.5 ± 0.1 | 3.6 ± 0.1 | 3.2 ± 0.1*, # | 3.07 ± 0.05*, # |
| AFP (ng/mL) | 0.8 ± 0.1 | 0.5 ±0.1 | 0.5 ±0.0 | 0.4 ± 0.1* |
| AST (U/L) | 97.4 ± 5.9 | 90.2 ± 4.8 | 196.8 ± 22.3*, # | 164.3 ± 13.9*, # |
| ALT (U/L) | 69.3 ± 3.9 | 71.2 ± 3.4 | 118.8 ± 16.7*, # | 94.3 ± 13.8 |
| ALP (U/L) | 199.2 ± 9.7 | 212.0 ± 7.6 | 164.8 ± 14.1# | 157.7 ± 1.6*, # |
| GGT (U/L) | 14.0 ± 3.0 | 15.3 ± 2.1 | 5.6 ± 2.1# | 2.5 ± 0.7*, # |
| PT (s) | 18.1 ± 0.5 | 19.2 ± 1.3 | 20.8 ± 0.6 | 19.4 ± 0.5 |
| TB (mg/dL) | 0.25 ± 0.04 | 0.2 ±0.0 | 0.2 ± 0.0 | 0.2 ±0.0* |
| Creatinine (mg/dL) | 0.4 ±0.0 | 0.3 ±0.0 | 0.3 ± 0.0 | 0.3± 0.0 |

| | | | | |
|---|---|---|---|---|
| Glucose (mg/dL) | 132.3 ± 7.8 | 141.1 ± 5.7 | 153.2 ± 7.7 | 148.2 ± 2.4 |
| Cholesterol (mg/dL) | 84.4 ± 4.3 | 86.2 ± 3.0 | 70.4 ± 3.8[*, #] | 64.8 ± 1.9[*, #] |
| Triglyceride (mg/dL) | 78.4 ± 11.1 | 70.3 ± 10.7 | 81.7 ± 21.0 | 112.1±15.2 |

**Supplementary Fig. 1. GNS561 is efficient in an orthotopic hepatocellular carcinoma patient-derived LI0752 xenograft BALB/c nude mouse model**.

Tumor volume (A) and tumor weight (B) in different groups: control, GNS561 at 30 and 50 mg/kg at day 41 after tumor inoculation. C, Serum alpha fetoprotein (AFP) at days 7 (D), 21 (E), 28 (F) and 41 (G) after tumor inoculation. Data represent the mean + SEM. Comparison of means was done by one-way ANOVA with Dunnett's post hoc analysis. *, $p < 0.05$.

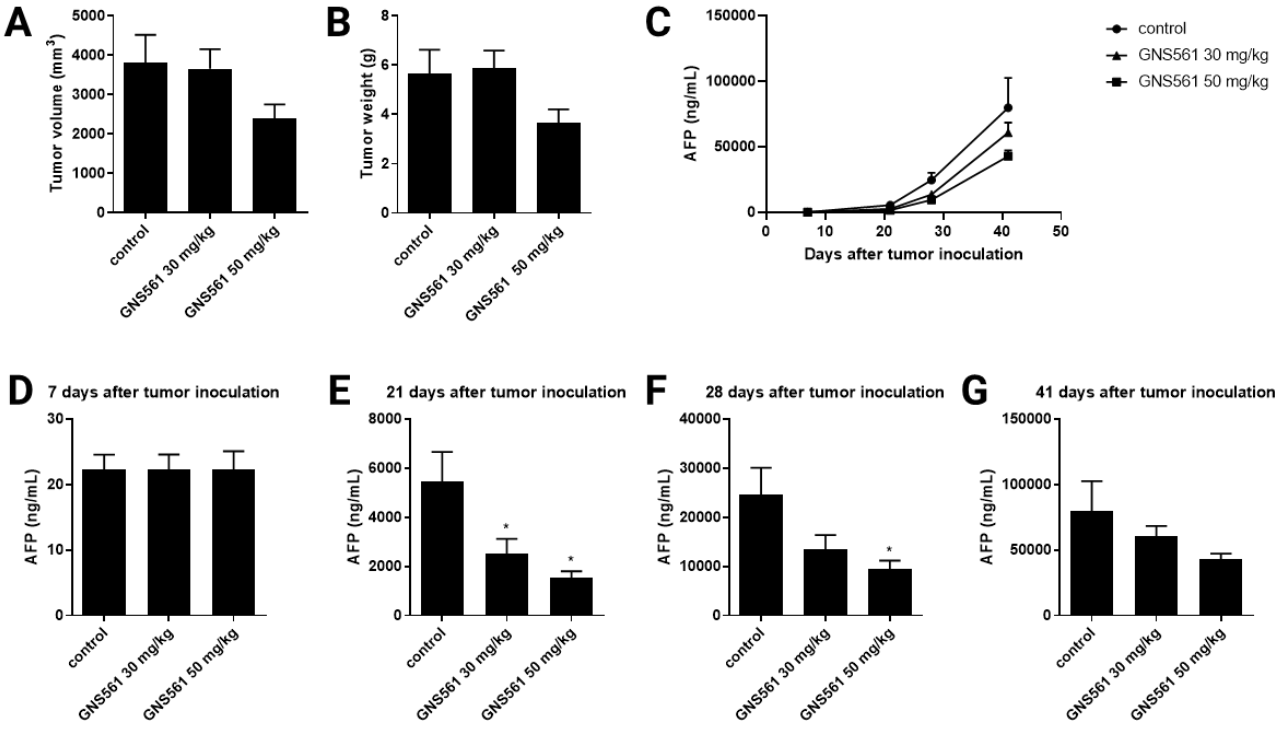

**Supplementary Fig. 2. Study design to assess the efficacy of GNS561 as single treatment and in combination with sorafenib in a diethylnitrosamine (DEN)-induced cirrhotic rat model of hepatocellular carcinoma.**

MRI (Magnetic Resonance Imaging)

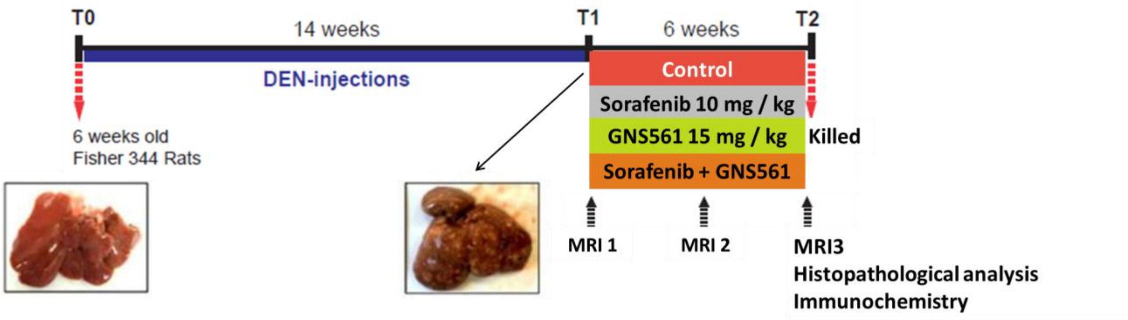

**Supplementary Fig. 3. GNS561 is a lysosomotropic agent**

Cell viability percent against vehicle condition after 24 h of treatment of HepG2 cells with GNS561 in the presence or absence of bafilomycin A1 (BafA1) (200 nM). Data represent the mean + SEM of three experiments. Student t-test was used for comparisons.*, $p < 0.05$.

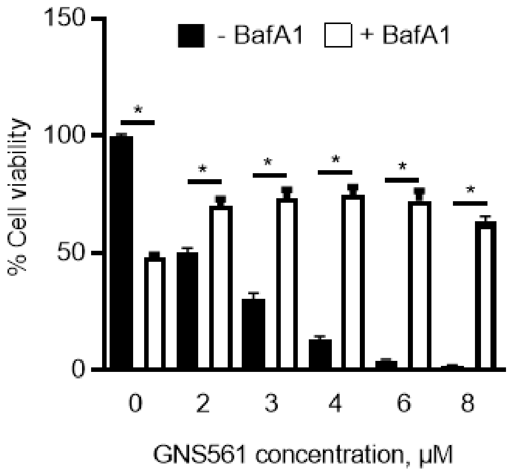

**Supplementary Fig. 4. GNS561 does not block directly enzymatic activity of cathepsins.**

Percent of peptidase activity of cathepsins B (CTSB), L (CTSL) and D (CTSD) in presence of GNS561 (4 and 10 µM) and E-64 (1 µM) or Pepstatin A (0.1 µM) calculated in comparison with the control condition. Data represent the mean + SEM of two experiments.

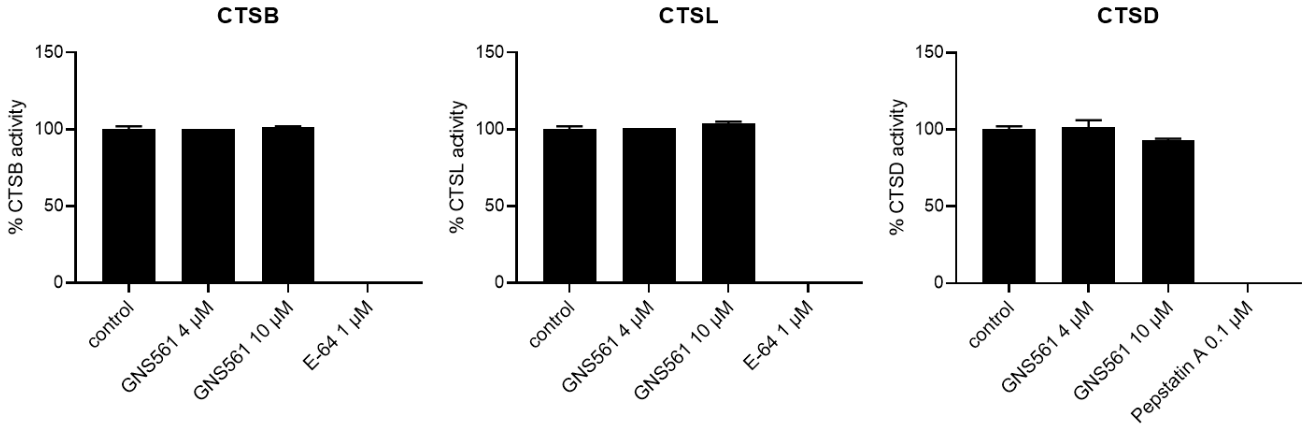

**Supplementary Fig. 5. GNS561 inhibits autophagy at late stage.**

Immunoblot analysis of light chain 3 phosphatidylethanolamine conjugate (LC3-II)blevels were performed in HepG2 (A) and Huh7 (B) cell lines incubated with vehicle (0 µM) or with indicated concentrations of GNS561 for 4 h or 24 h in the presence or absence of bafilomycin A1 (BafA1) (100 nM, 2 h). Glyceraldehyde-3-phosphate dehydrogenase (GAPDH) was used as a loading control. As indicated under each lane, the autophagic flux, determined as the ratio between the LC3-II level normalized against GAPDH level (Norm LC3-II) with BafA1 and without BafA1 (No BafA1), is expressed in arbitrary units. Three independent experiments were performed. Representative autoradiograms are shown.

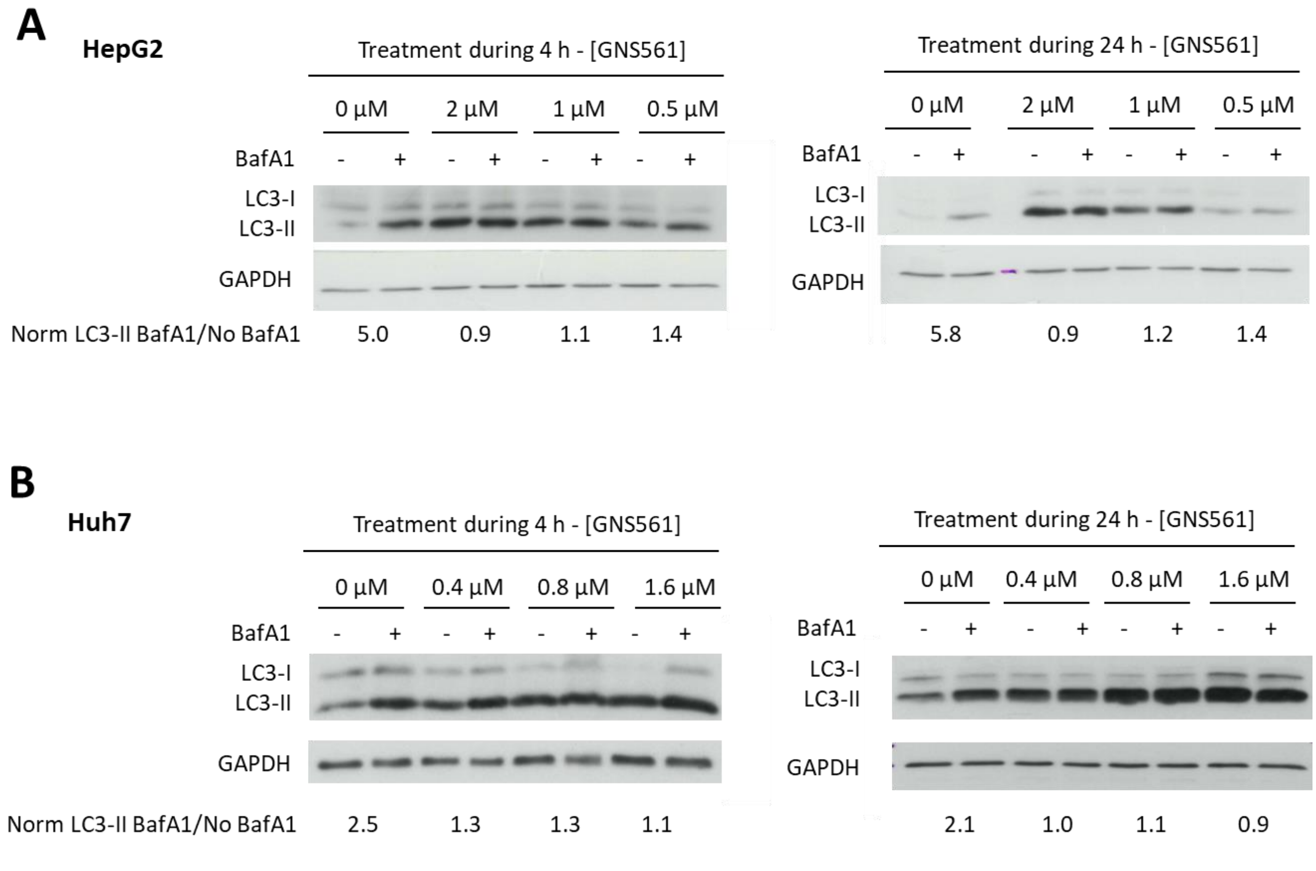

**Supplementary Fig. 6. NtBuHA partially rescues the anti-tumor activity of HCQ.** Cell viability percent against vehicle condition after 24 h of treatment with HCQ in the presence or absence of N-tert-butylhydroxylamine (NtBuHA) (8 mM). Data represent the mean + SEM of three experiments. For comparison, Student t-test was used. *, $p < 0.05$.

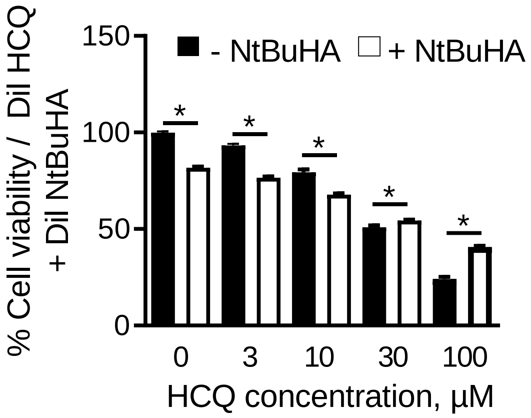

**Supplementary Fig. 7. NtBuHA doesn't affect GNS561 lysosomal localization**

Pearson correlation coefficient between GNS561G (GNS561 analog tagged with Bodipy FL) and lysosomal-associated membrane protein 2 (LAMP2) after treatment with GNS561 for 2 h in the presence or absence of $NH_4Cl$ (20 mM) or N-tert-butylhydroxylamine (NtBuHA) (8 mM). Three independent experiments were performed. For comparison with GNS561 alone, one-way ANOVA with Dunnett's post hoc analysis was used.*, $p < 0.05$. Box and whisker representation (min to max) is shown.

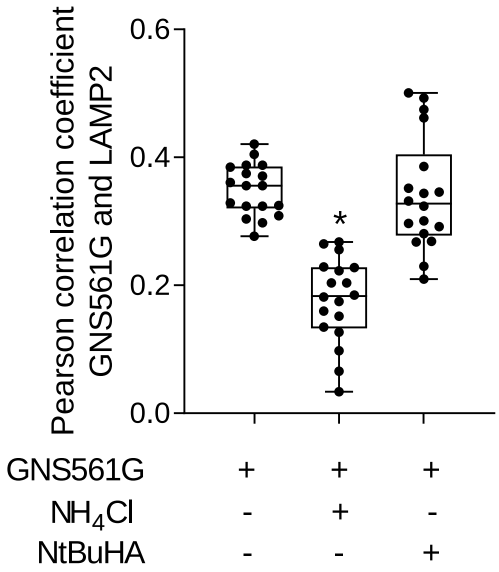

**References**

1. Bestion E, Jilkova ZM, Mège J-L, Novello M, Kurma K, Pour STA, et al. GNS561 acts as a potent anti-fibrotic and pro-fibrolytic agent in liver fibrosis through TGF-β1 inhibition. Therapeutic Advances in Chronic Disease 2020;11:2040622320942042.

2. Galibert M, Wartenberg M, Lecaille F, Saidi A, Mavel S, Joulin-Giet A, et al. Substrate-derived triazolo- and azapeptides as inhibitors of cathepsins K and S. European Journal of Medicinal Chemistry 2018;144:201-210.

3. Wartenberg M, Saidi A, Galibert M, Joulin-Giet A, Burlaud-Gaillard J, Lecaille F, et al. Imaging of extracellular cathepsin S activity by a selective near infrared fluorescence substrate-based probe. Biochimie 2019;166:84-93.

4. Brun S, Bassissi F, Serdjebi C, Novello M, Tracz J, Autelitano F, et al. GNS561, a new lysosomotropic small molecule, for the treatment of intrahepatic cholangiocarcinoma. Investigational New Drugs 2019;37:1135-1145.

5. van Diggelen OP, Keulemans JL, Winchester B, Hofman IL, Vanhanen SL, Santavuori P, et al. A rapid fluorogenic palmitoyl-protein thioesterase assay: pre- and postnatal diagnosis of INCL. Molecular Genetics and Metabolism 1999;66:240-244.

6. Jilkova ZM, Kuyucu AZ, Kurma K, Ahmad Pour ST, Roth GS, Abbadessa G, et al. Combination of AKT inhibitor ARQ 092 and sorafenib potentiates inhibition of tumor progression in cirrhotic rat model of hepatocellular carcinoma. Oncotarget 2018;9:11145-11158.
